# Active Conformations of Neuronal Na^+^,K^+^-ATPase isoforms and a Disease-Causing Mutant

**DOI:** 10.1101/2025.07.27.666930

**Authors:** Mads Eskesen Christensen, Michael Habeck, Adriana Katz, Marlene Uglebjerg Fruergaard, Yoav Peleg, Uri Pick, Steven J D Karlish, Poul Nissen

**Author notes:** these authors contributed equally (Mads Eskesen Christensen, Michael Habeck).

## Abstract

Na^+^,K^+^-ATPases establish and maintain the vital electrochemical gradients for Na^+^ and K^+^ across animal cell membranes. The protein is a ternary complex composed of α, β and FXYD subunits, of which isoforms that fine-tune transport properties are expressed in a tissue-specific fashion. Here we report cryo-EM structures under active ATPase turn-over conditions of the ubiquitously expressed human α1β1FXYD1 and neuron-specific α3β1FXYD1 isoform complexes and probe their specific functional and biophysical properties. The data provides an extensive insight into Na^+^-transport of ATP-activated enzyme through four distinct conformational states, including a long-sought sodium-bound phosphoenzyme intermediate, denoted [Na_3_]E2P. This hitherto elusive conformation reveals a crucial structural change that precedes Na^+^ release in the inward to outward (E1P-E2P) transition, within the general context of the sequential, active transport mechanism. We investigated and discuss the mechanism of the physiologically important differentiation in Na^+^ affinity of α3 compared to α1, the co-operative Na^+^ binding at the ion-binding sites, and the mechanistic aspects of cytoplasmic ion gating and extracellular Na^+^ release. Finally we present the first structures of a disease-causing mutant form of α3, associated with Alternating Hemiplegia of Childhood (Q140L). The mutation compromises a specific phospholipid-binding pocket and impedes polyunsaturated phospholipid-mediated stimulation of Na^+^,K^+^-ATPase activity.

## Introduction

Na^+^,K^+^-ATPases maintain the uneven distribution and electrochemical gradients for Na^+^ and K^+^ ions across the plasma membrane of animal cells. The ATP-driven cycle of active Na^+^ and K^+^ transport follows the Albers-Post-mechanism, where the enzyme alternates between inward- and outward-oriented conformations denoted E1 and E2, respectively (**see Suppl. Fig. 1** for detail) [1–3]. The transport cycle is overall electrogenic, resulting in the extrusion of three cytoplasmic Na^+^ in exchange for two extracellular K^+^ ions [4]. The resulting ion gradients and derived membrane potential fuel a range of physiologically vital processes, including ion channel signalling, cell volume control, and secondary active transport of ions, nutrients and metabolites.

The Na^+^,K^+^-ATPase is an obligate hetero-oligomer consisting of the catalytic α, the accessory β, and a regulatory FXYD subunit. Isoforms of the three subunits are encoded by different genes in mammals, specifically ATP1A1 through 4, ATP1B1 through 3, and FXYD1 through 7 for α, β and FXYD subunits, respectively. Different isoform combinations exhibit distinct kinetic properties and ion affinities, adapted for specific cellular properties and functions [5–7]. The α1β1 complex is ubiquitously expressed and considered the housekeeping form, whereas α3 is expressed in neurons, primarily with β1 or β3, and displays a reduced apparent affinity for cytoplasmic Na^+^ ions as compared to α1 [8].

Neuronal α3 complexes play critical roles in brain physiology, acting as auxiliary pumps in parallel with the ubiquitous α1. Following neuronal activity, intracellular Na^+^ concentrations increase, and α3 complexes become transiently activated, thus aiding the rapid restoration of cytoplasmic Na^+^ to basal levels [8]. Rare dominant mutant forms of α3 cause brain diseases, classified as *ATP1A3* related disorders [9, 10]. Different sites and types of neuronal α3 mutations manifest different neurological disorders, in particular Rapid-onset Dystonia with Parkinsonism (RDP) and the severe Alternating Hemiplegia of Childhood (AHC) [11]. RDP mutations are widely distributed across the *ATP1A3* gene and generally show loss-of-function, suggestive of different mechanisms of inactivation and haploinsufficiency. By contrast, AHC mutations generally appear as missense mutations and cluster mostly within or close to the transmembrane domain, potentially associated with more specific mechanisms [12]. An understanding of AHC mechanisms could provide insights into ways of circumventing AHC-specific pathology.

Functional integrity of Na^+^,K^+^-ATPase is tightly coupled to the lipid composition of the membrane. Indeed several selective interactions of phospholipids with the protein in specific binding pockets have been described. These protein-phospholipid interactions either stabilize, stimulate or inhibit Na^+^,K^+^-ATPase activity, respectively [13, 14].

Whereas crystal structures of pig kidney and shark rectal gland α1 isoform complexes and, most recently, also cryo-EM structures of mammalian complexes have shed light on individual states stabilized by inhibitors or mutations [15–17], information on the human Na^+^,K^+^-ATPase isoform complexes with full functional integrity and determined in active turn-over conditions, have been lacking. Furthermore, structure-based insight into Na+ release, isoform specific properties, and mechanisms of disease-causing mutations remain elusive. Here, we present structural and functional studies of human wild-type α1β1FXYD1 and α3β1FXYD1 complexes, and an AHC disease-causing α3 mutation (Q140L) that provide profound, new insights into these questions.

## Results and Discussion

### Cryo-EM structures of ATP-activated Na^+^,K^+^-ATPases

Human Na^+^,K^+^-ATPase complexes of α1- and α3-isoforms with His_10_β1-subunit, were expressed in *Komagataella phaffii (Pichia pastoris)*, purified by affinity chromatography and reconstituted with FXYD1 and lipid mixtures in saposin A lipoprotein nanodiscs [18]. The resulting nanoparticles were subsequently purified by size exclusion chromatography. **(Suppl. Fig. 2)**. Cryo-EM data of α3β1FXYD1 and α1β1FXYD1 complexes were obtained in the presence of Na^+^, ATP and Mg^2+^, conditions in which the Na^+^,K^+^-ATPase performs Na^+^-transport cycles [19]. For both complexes we identified four distinct conformations in the data sets **(Fig. 1A, 2A)** associated with Na^+^ transport: i) a pre-phosphorylated Na_2_E1-ATP state, ii) a sodium-occluded [Na_3_]E1P-ADP complex formed by autophosphorylation, iii) a hitherto unknown Na^+^ occluded phosphoenzyme intermediate, denoted [Na_3_]E2P, and iv) an outward-open, Na^+^ released E2P state. The structures provide a near complete view of Na^+^ transport, unbiased by stabilizing mutations, detergents or inhibitors and revealing the sequential steps of sodium binding, phosphorylation, E1P-E2P transition and ion-release.

**Figure 1.**
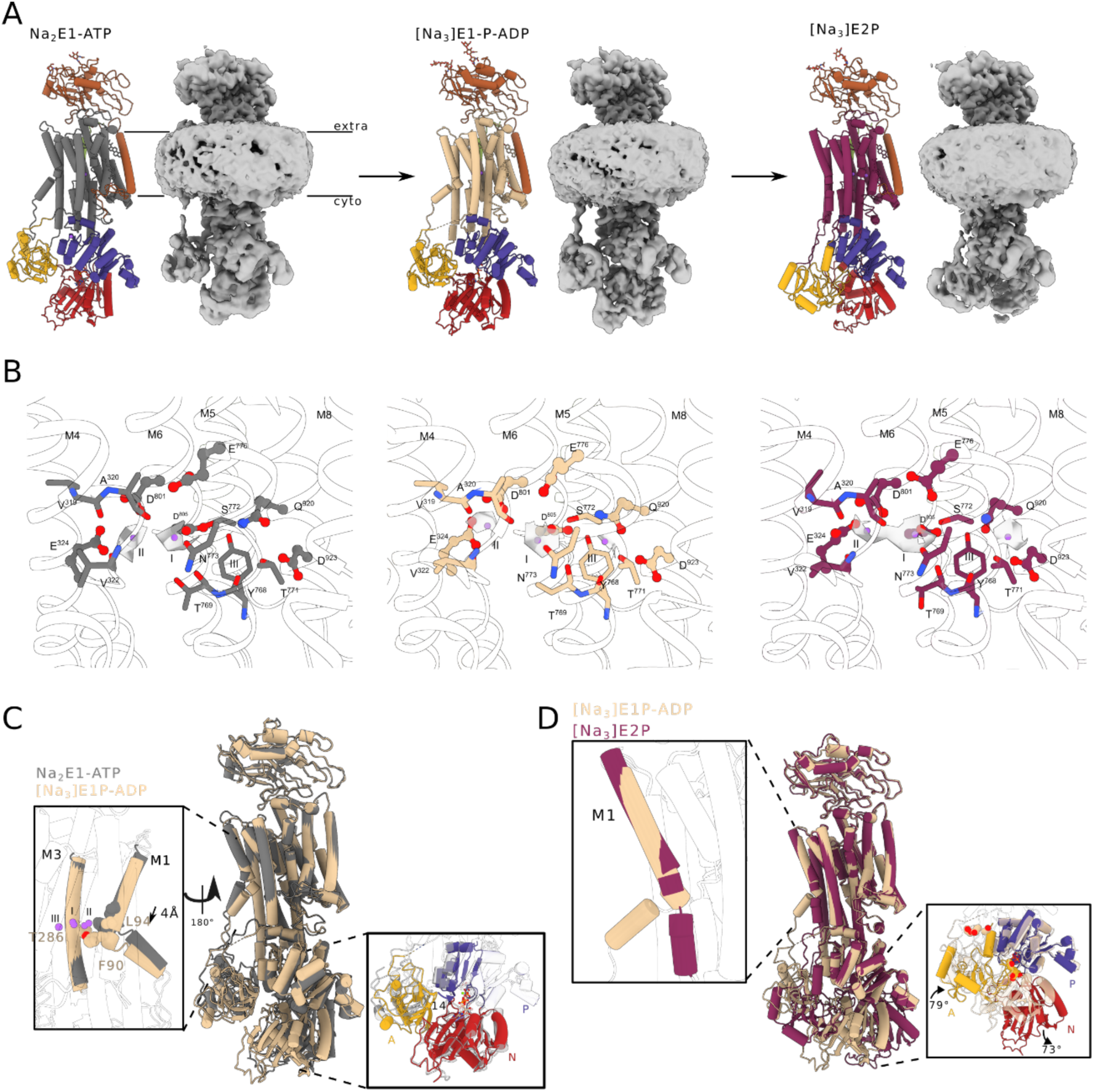
Structures of Na^+^-bound states of α3-Na,K-ATPase. (A) Sodium-bound cryo-EM structures of wt α3ß1FXYD1 Na,K-ATPase with α-A domain in yellow, α-P domain in blue, α-N domain in red, ß-subunit in orange, FXYD1 in light green (barely visible in the back), and the α-TM-domain in grey, wheat and maroon, respectively, for the three indicated states. (B) cryo-EM density around the Na^+^ binding sites I, II and III of the respective states shown in A (contoured at 4 s.d. for Na_2_E1-ATP and [Na_3_]E1P-ADP, and at 3.5 s.d. for [Na_3_]E2P). Site III only shows density for a bound ion in the occluded [Na_3_]E1P-ADP and [Na_3_]E2P states. Hence, we propose a binding mechanism occupying first exposed binding sites I and II, then with internal reorganization leading to occupation of site III, co-operative binding of a third Na+ at exposed sites I/II and occlusion/phosphorylation. In [Na_3_]E2P, site III is shifted towards M8 with D923. We propose that the shift breaks the co-operativity of the three sites and initiates extracellular release. (C) Alignment of Na_2_E1-ATP and occluded [Na_3_]E1P-ADP states showing the closure of the cytoplasmic gate by the movement of M1. The cytoplasmic domains line up for phosphoryl transfer. (D) Alignment of Na^+^-occluded [Na_3_]E1P-ADP and [Na_3_]E2P states. Rotation of the A-domain relative to the P-domain as well as the rotation of the N-domain favour ADP release, which essentially becomes an irreversible step that seals the E1P-to-E2P transition. Straightening of M1 is the major change in the transmembrane domain and is coupled to the reconfiguration of the cytoplasmic domains.

**Figure 2.**
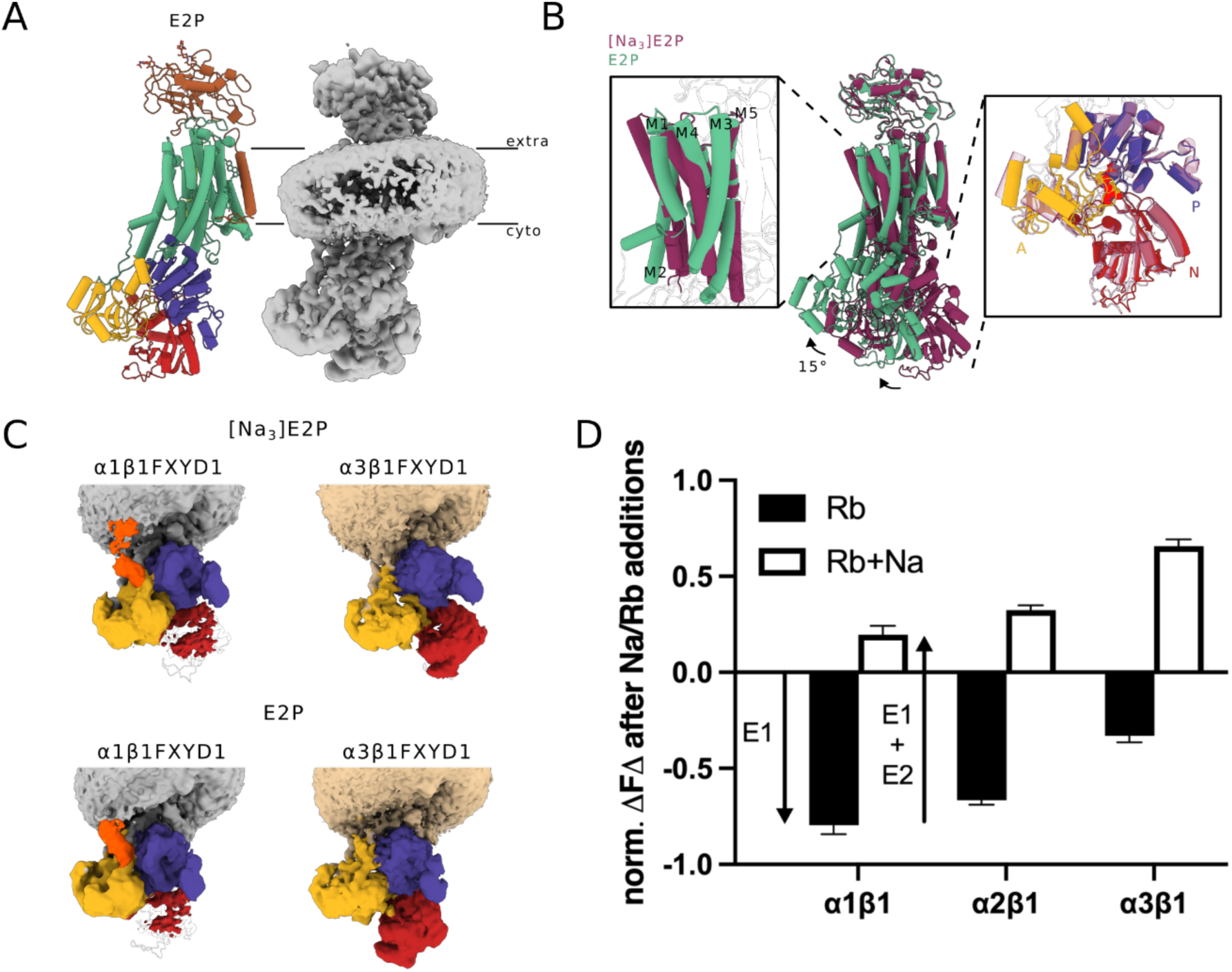
E2P states of α1 and α3 Na,K-ATPase. (A) Structure and cryoEM map of wt α3ß1FXYD1-Na,K-ATPase. Colours as in Fig. 1 except the α-TM domain now in green. (B) Major rearrangements of the [Na_3_]E2P to E2P transition showing the rearrangement of TM1-5 and alignment of the cytoplasmic domains, which rotate in a coordinated 15° angle. (C) Density of the cytoplasmic domains of wt α1 and α3 isoforms in the [Na_3_]E2P versus outward open E2P states at a contour level of 2.5. The density likely representing the far N-terminus of the A-domain, which extends towards the saposinA nanodisc, is highlighted in orange. Other colours as in A. This density is only visible in α1 but not α3, while the N-domain is better defined α3 versus α1. (D) E1/E2 equilibria of FITC-labelled Na,K-ATPase isoforms estimated from amplitude of fluorescence changes upon addition of 20mM RbCl and then 100mM NaCl. The data represent averages of 3 to 5 determinations ± SEM. Values of E1/(E1+E2) in the Na− and K-free high ionic strength medium are as follows: α1β1 0.86±0.1; α2β1 0.67±0.02; α3β1 0.33±0.03.

These structures at turn-over were determined to an overall resolution of 2.9-3.3 Å for the neuronal α3β1-FXYD1 (GSFSC 0.14) and 3.2-3.5 Å for the ubiquitously expressed α1β1-FXYD1 (**Supplementary table 1**). Hence, mechanistic features shared between isoforms are primarily derived from the α3β1-FXYD1 data and, generally, we refer to human α3 residue numbers unless otherwise stated.

We also obtained full turnover data for an α3β1-FXYD1 sample with both Na^+^ and K^+^ present, yielding similar conformations (except for E2P being replaced by [K_2_]E2.P, see further below), but at reduced quality, most likely due to the increased Na^+^,K^+^-ATPase turnover rate and flexibility with saturating K^+^ concentrations **(Suppl. Fig. 3)**.

### The cytoplasmic sodium-gating mechanism

For the inward-open Na_2_E1-ATP conformation, an open pathway between M1, M3 and M4 extending from the cytoplasm to the ion-binding sites is observed **(Suppl. Fig. 4)**. Na^+^ sites I and II appear ordered with density for bound ions, while density likely representing ATP is observed in the more flexible N-domain. Notably, Na^+^ site III appears vacant in this state and the conformation most likely represents a pre-phosphorylated intermediate prior to Na^+^ binding at site III, indicating it as a rate limiting step. For the occluded [Na_3_]E1P-ADP state, density is observed for all three Na^+^ sites, and a bipyramidal arrangement reveals the phosphoryl transfer from ATP to Asp366, similar to earlier structures for ADP-AlF_x_ stabilized forms [17, 20, 21].

Kanai et al. have previously suggested a serial site III-I-II binding of the three Na^+^ ions in the E1-ATP to Na3E1-ATP transition that activates phosphorylation [20]. Our results however, indicate that sites I and II are occupied first **(Fig. 1B)**, with subsequent Na^+^-binding at the deeper site III. Using two fluorescent reporters monitoring changes in cytoplasmic and transmembrane domains, Schneeberger & Apell have shown that binding of the first two Na^+^ ions is not electrogenic [22], consistent with Na^+^-binding to the accessible sites I and II through the solvated pathway observed in the Na_2_E1-ATP state. Subtle conformational changes, possibly coupled to deprotonation and the interplay with a proximal site IIIa [21], may allow an internal push-through mechanism for transfer of a Na^+^-ion to site III, with positive cooperativity for binding of a third Na^+^ ion via the exposed sites I and II [23].

Comparing the Na_2_E1-ATP and [Na_3_]E1P-ADP states, the TM domains appear rather similar, except for a 4 Å shift of M1 towards the cytoplasmic interface (Fig 1C). Upon Na^+^ binding at site III, the shift of M1 closes a local perturbation of the membrane interface, occluding the bound Na^+^-ions and stabilizing a configuration of the cytoplasmic domains that promotes phosphorylation (**Fig 1C**). Binding of the third Na^+^ ion has been shown to involve subtle conformational changes [22], and an intramolecular, electrogenic transfer of Na^+^ into a neutral site III was proposed (which could be the internal push-through mechanism proposed above). Similarly to the observed 4Å shift of M1 along the TM-domain, the gating of SERCA is described as a ‘sliding door’ mechanism of M1, although with a larger ∼12Å shift relative to M4 [24]. The observed shift of M1 and the ‘sliding door’ ion-occlusion mechanism is in agreement with a recent cryo-EM structure of ATPγS bound human Na^+^,K^+^-ATPase (α_1_β_1_FXYD2) [25]. However, we do not find any support for a so-called ‘hinged door’ gating mechanism, in which M1 tilts within the plane of the membrane, as proposed in a cryo-EM study of human α_3_β_1_FXYD6 stabilized by the non-hydrolyzable ATP analog AMPPCP and inactivating mutations at the ion binding sites [17].

Interestingly, residues within site III show no compelling local differences upon Na^+^ binding. In comparison, Ca^2+−^ATPases feature only sites I and II, or even site II exclusively for activation of phosphorylation. So, what causes occupation of site III in the Na^+^,K^+^-ATPase to trigger Na^+^-dependent phosphorylation? We propose that the cooperativity of all three sites is the key. Only upon binding of Na^+^ at site III, will sites I and II gain the kinetic stability to support the stable interactions with M1/M4 and induce the conformational changes associated with phosphorylation. Indeed, mutation of a conserved glutamate in M4 (Glu324 for α3) at site II causes reduced Na^+^-affinity, decreased phosphorylation and destabilised occlusion [26, 27], while mutation of residues equivalent to Asp801 and Asp805 bridging between Na^+^-sites I, II and III, presumably critical for cooperativity, exhibit pleiotropic effects on Na^+^,K^+^-ATPase function [28]. Cooperativity of the three Na^+^ sites ensures that the pump moves forward through the catalytic cycle only upon binding of all three Na^+^ ions, despite a relatively low Na^+^-affinity and competition with K^+^ ions for sites I and II.

### Sodium release, the [Na_3_]E1-P-ADP → [Na_3_]E2P → E2P reactions

Following phosphorylation and ADP release, the enzyme undergoes a transition that leads to the release of sodium to the extracellular side. Intermediate states associated with this critical reaction have been elusive, but here at turn-over conditions, a hitherto undetected intermediate is determined at 3.1 Å overall cryo-EM map resolution. Except for M1 and site III (see further below), the TM domain largely resembles that of the [Na_3_]E1P-ADP state. By contrast, the cytoplasmic domains have reconfigured to a nucleotide-free E2P-like form (**Fig. 1D, Suppl. Fig. 4B**), in which the A-domain with the TGES loop (catalyzing dephosphorylation in a next step [15, 29, 30]) interacts with and shields the phosphorylated Asp366 at the P-domain. Sites I and II are similar with density consistent with Na^+^ binding, but interestingly, site III appears shifted by about 2.2 Å towards M8 (**Fig 1B**), moving away from Thr771 and now appearing as additional density located adjacent to Asp923 and Asn920. The assignment of a shifted Na^+^ site III is tentative at the given resolution, but other models seem less obvious, and we therefore denote this conformation [Na_3_]E2P, which distinguishes it from [Na_3_]E1P and E2P states.

Relative to [Na_3_]E1P-ADP, when superimposed on the TM domain, the cytoplasmic headpiece of [Na_3_]E2P is rotated and reconfigured with large conformational changes. This includes a 80° rotation of the A-domain towards the phosphorylated residue of the P-domain (see above). In turn, this forces M1 into a unique, lateral configuration parallel to M2-M4 (**Fig. 1D**) with the linker between the A-domain and M1 (A-M1) adopting a stretched conformation. The empty nucleotide site at the N-domain is exposed and is now entirely detached from the phosphorylation site, displaying a defining feature of E2P forms as an ADP-insensitive phosphoenzyme.

Although this intermediate state ([Na_3_]E2P) has not been observed previously, a similar calcium-occluded E2P state for SERCA has been inferred from biochemical studies [31, 32], MD simulations [33], and from x-ray solution scattering studies [34]. Single-molecule FRET studies [35] and a recent cryo-EM structure of the bacterial Ca^2+^-ATPase LMCA1 carrying a four-glycine insert in the A-M1 linker [36] reveal a similar conformation. Thus, these studies of the homologous Ca^2+^-ATPases are in agreement with the [Na_3_]E2P structure presented here regarding the overall conformation, with an E2P configuration of the cytoplasmic domains and the E1P-like conformation of the TM-domain, imposing strain on the A-M1 linker in the E1P-E2P transition.

In support of preserved Na^+^ sites I and II and a shifted site III for [Na_3_]E2P, charge-pulse experiments have indicated that the [Na_3_]E1P-ADP → [Na_3_]E2P transition exhibits only a minor electrogenic coefficient, minor movements of the Na^+^-ions, and small rearrangements of charged residues in the membrane [37]. Following this model, the shift of Na^+^-site III towards M8 may account for the minor movement that subsequently destabilizes the other ion binding sites (due to loss of cooperativity, see below), causing a rearrangement of TM1-4 and the formation of the extracellular exit pathway. The role of site III Asp923 and Asn920 on Na^+^ binding has been investigated previously [38, 39]. While mutations of Asp923 reduce the Na^+^ affinity on the intracellular as well as the extracellular side, mutation of Asn920 predominantly affects Na^+^ binding from the extracellular side. Nielsen et al. [35] point out that the Na^+^ ion in site III may have some freedom to move around. Thr771, which is associated with the shift of site III, also shows effects on Na^+^ dependent activity when mutated to Alanine [40] and mutation of nearby Pro775 to Leu is associated with milder ATP1A3 disorders [41].

Electrophysiological measurements on squid giant axons have revealed further details of a sequential sodium release process from Na,K-ATPase with a slow first step exhibiting a majority of the charge component, interpreted as deocclusion and release of the first Na^+^ through a narrow pathway. This is followed by a medium and fast component, which carry only a small fraction of the electric current, inferred as release of the second and third Na^+^ ion through a wider, solvated channel [42, 43].

We speculate that the small shift of site III breaks cooperativity in the [Na_3_]E1P-ADP to [Na_3_]E2P transition and initiates the release of Na^+^. MD-simulations suggest that the change of protonation of D801, D805 and D923 between E1P and E2P is crucial [44]. In particular, protonation of D923 was suggested to induce Na^+^ release [45]. Razavi et al. proposed a model in which an anion binds to the cytoplasmic M8-9 loop and triggers the change in protonation [46]. In this model D923 becomes protonated in E2P from a cytoplasmic access channel [47], while E776 donates a proton to D805 and D801 to E324. Our maps do not show density for an anion at the suggested cytoplasmic site, but these simulations are overall in agreement with the destabilization of Na^+^ sites through a site III shift observed here. Following this initial step, release from site II, likely through a narrow pathway between M1-M2, M4 and M6, may trigger the opening and further release from sites I and III through the fully solvated pathway observed in the outward-open E2P state (**see Suppl. Fig. 5 for a proposed Na^+^ binding and release sequence**).

Finally, the outward-open E2P conformation observed here is similar to published crystal structures of E2-BeF_x_ and E2P-ouabain [15, 30, 48]. Superimposing the ion-occluded [Na_3_]E2P intermediate on αM7-10 of E2P (a rigid subdomain of the TM domain) shows significant conformational differences in αM1-6 of the TM domain, and a 15° rotation of the cytoplasmic headpiece, which moves essentially as a rigid body (**Fig. 2B**). The conformational changes result from the release of Na^+^ and relaxes at the same time the extended M1 and stretched A-M1 linker associated with the [Na_3_]E2P state.

The structural characterization of an E1P-E2P intermediate state of Na^+^,K^+^-ATPase ([Na_3_]E2P) points to a sequential E1P to E2P transition mechanism, in which the initial response to phosphorylation is the rearrangement of the cytoplasmic domains. The A domain rotates and comes into close proximity with the P domain, protecting the active site aspartyl-phosphate from spontaneous dephosphorylation and futile cycling without transport. In addition, the [Na_3_]E2P conformation provides structural evidence for ADP release preceding the E1P-E2P transition and Na^+^ deocclusion. The structure highlights the importance of TM1 and the A-domain linker(s) as key components in the transmission of conformational rearrangements, linking the initial reconfiguration of the cytoplasmic domains to a subsequent rigid body movement of the cytoplasmic headpiece and rearrangement of the transmembrane helices associated with Na^+^ release. Indeed, insertion of additional residues in the A-M1 linker of related SERCA and LMCA1 relaxes strain and reduces allosteric coupling between the cytoplasmic and TM-domain, pausing the de-occlusion and release of calcium [31, 32, 36].

As mentioned earlier, we also recorded a turn-over dataset with the addition of 20mM KCl, i.e. stimulating the full transport cycle. Overall, it shows less ordered structures in the cryo-EM analysis, but we observe also here the (Na/K)_2_E1-ATP, [Na_3_]E1P-ADP and [Na_3_]E2P conformations, as well as a [K_2_]E2.P conformation (**Suppl. Fig. 3**) instead of the outward-open E2P. This latter state is expected, since K^+^ binding is fast and shows only little competition with Na^+^ to the E2P conformation. Hence the limiting steps under full cycle turn-over conditions applied here appear to be i) binding of the third Na^+^ ion, ii) autophosphorylation and ADP-release, iii) Na+ release from [Na_3_]E2P, and iv) dephosphorylation and P_i_ release leading into cytoplasmic K^+^-release.

### Structural Comparison and Na^+^ affinity of α1 versus α3 isoforms

Overall, the structures of α3 and α1 are similar (**Suppl. Fig. 6**), however, with one significant structural difference. Whereas the α3-maps show clear features for all three cytoplasmic domains for all states, the maps for α1 complexes are consistently weak for the N-domain in all but the [Na_3_]E1P-ADP state (**Fig. 2C**). This indicates a far more flexible N-domain for α1 as compared to α3. By contrast, α1 maps show more clearly the extended features for the N-terminal tail reaching the membrane interface of the saposin nanodisc, in particular for [Na_3_]E2P and outward-open E2P (**Fig. 2C**). Indeed, several α1- and α3-specific residues are located at the N-terminal tail, which is partially disordered and 10 residues shorter in α3, when compared to α1. As discussed below, these structural features may be relevant to functional differences.

The major reported differences in functional properties of the housekeeping α1 and neuronal α3 isoforms are the lower apparent affinity for cytoplasmic Na^+^ and a somewhat lower voltage dependence of α3 [8, 49]. A lower Na^+^ affinity could reflect intrinsic differences in the Na^+^ binding sites, but we observe no prominent differences for the highly conserved Na^+^ binding residues of α1 and α3, respectively. Alternatively, a difference in E1/E2 conformational equilibrium of α3 versus α1 might be responsible for the difference in apparent Na^+^ affinity. We tested this concept using fluorescein isothiocyanate (FITC), a fluorescence probe of E1-E2 conformations (**Fig. 2D**) [50], which selectively labels a lysine sidechain at the nucleotide binding site [51]. In a Na^+^ and K^+^-free buffer at high ionic strength, a condition known to strongly stabilise E1 [52], the addition of Rb^+^ (a congener of K^+^) induces the transition to rubidium-bound E2 that quenches fluorescence, while subsequent addition of Na^+^ reverses the fluorescence change upon return to the sodium-bound E1 conformation. As seen in **Fig.2D** for α1, addition of 20mM Rb^+^ in the high ionic strength medium lacking both Na^+^ and K^+^ ions, induced fluorescence quenching, which reverted to slightly above the starting level upon addition of 100mM Na^+^. This confirms that α1 is indeed largely stabilized in the E1 conformation in the initial condition. For α3, by contrast, addition of 20mM Rb^+^ induced a much lower fluorescence quenching, but a significant increase above basal level upon addition of 100mM Na^+^, showing that α3 is predominantly in the E2 conformation in the initial condition^1^. In a cellular environment, a conformation of α3 poised towards E2 will result in a stronger competition of cytoplasmic K^+^ for binding of Na^+^ ions and, thus, a lower apparent Na^+^-affinity of α3 compared to α1.

A hypothesis for the distinct conformational preferences of α3 and α1 relates to the observed differences in the N domain flexibility. Indeed, the majority of unique α3-residues are located in the A and N-domains, which interact in the E2 conformation. Thus, for α3, the E2 conformation may be stabilized by stronger interactions of a more ordered N-domain with the A domain. Crystal structures of pig and bovine kidney α1β1 enzyme in E2P-like conformations typically show an ordered N-domain [30, 53, 54], but it adopts different positions depending on crystal packing indicating again flexibility. Recent cryo-EM structures of shark α1β1 show reduced local resolution for the N-domain as well (and A-domain) [13], although less pronounced than seen here for ATP activated human α1β1.

The N-terminal segments of α1 and α3 also show differential interactions. The density of the α1 N-terminal segment approaching the saposin lipoprotein nanodisc hints at a functional role at the membrane interface. Indeed, studies by limited proteolysis and mutations revealed that the N-terminal tail of α1β1 Na^+^,K^+^-ATPase affects the E1-E2 conformational equilibrium [55, 56]. More recently, the interaction of the lysine-rich N-terminal tail with the membrane was proposed as a regulator of the conformational equilibrium [57, 58]. It is puzzling, however, that both E2 and E1 stabilization have been proposed. Hossain et al. speculated that the degree of disorder and transition of the tail from a structured to a disordered state could stabilise the E1 conformation [59]. On the other hand, chimeras of α1/α3 replacing also the N-terminal segment did not show significant changes in Na^+^ affinity [60]. Potentially, interaction of the N-terminal segment with the membrane might provide a mechanism for voltage-dependent regulation of transport rates.

All considered, no individual residues or segments can be pinpointed as a root cause of the lower Na^+^ affinity of α3. Rather, many small contributions of α3-specific residues could contribute to a more stable configuration of domains in E2 states, most notably for the N-domain, and thereby a reduction in apparent Na^+^-affinity.

### Structure and functional properties of an AHC-causing mutation

More than 150 different rare mutations of *ATP1A3* causing AHC have been described [61]. We present here the first structures of an AHC-causing mutant of α3 (Q140L), which causes a severe AHC phenotype [10, 12]. Q140 is located at a specific phospholipid site between αM2, 4, 6, 9 and FXYD1, termed site B [14]. The site is associated with the binding of poly-unsaturated phosphatidylethanolamine or phosphatidylcholine (e.g. 18:0,22:6 PE), which stimulate Na^+^,K^+^-ATPase activity [14, 62]. The α3-Q140L, the equivalent Q150L mutation of α1, and α1-Q150H (equivalent of α3-Q140H, which is associated with a less severe disease phenotype, termed D-DEMØ [63]), were expressed like WT complexes. For initial characterization of WT versus the mutants, the Na^+^,K^+^-ATPase activity and turn-over rates were determined in yeast membranes. Both α3-Q140L and the equivalent α1-Q150L showed strongly reduced, although not fully inactivated Na^+^,K^+^-ATPase activity and turnover rates (< 20% of WT, **Suppl. Table 2**). By contrast, the D-DEMØ equivalent α1-Q150H showed only moderately reduced Na^+^,K^+^-ATPase activity, which may be of interest in correlation to a milder disease phenotype.

Unlike several other AHC mutant forms of α3, the α3-Q140L protein can be purified making it amenable to structural studies. Cryo-EM structures for the α3-Q140L mutant were then determined under the Na^+^ transport conditions also used for wild-type α3. The same four conformations were detected as for WT α3 with rmsd of 0.85-1.22Å between the respective sub-class structures, and generally with only small differences in the arrangement of the cytoplasmic domains relative to the TM domain (**Fig. 3A**). While density for cholesterol sites appear similar between WT and the mutant structures (not shown), important differences were observed for lipid binding site B (**Fig. 3B**). The maps of WT α3 show clear density for a lipid at site B in both [Na_3_]E1P-ADP and [Na_3_]E2P conformations, but the maps for α3-Q140L show only weak density for a bound lipid, if present at all (**Fig. 3B**).

**Figure 3.**
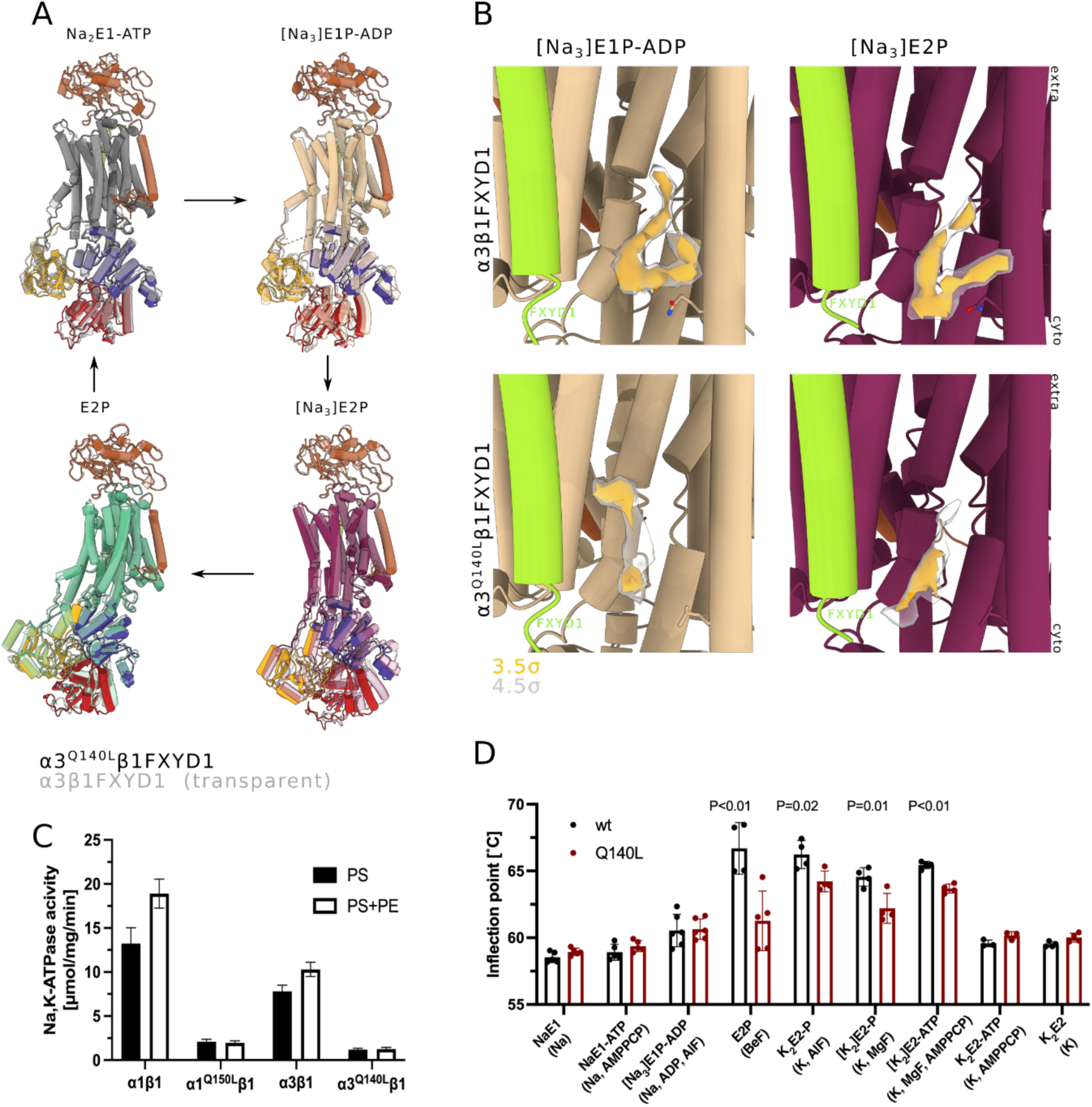
Characterization of the Q140L AHC variant. (A) Alignment of wt α3 and α3-Q140L showing overall similar structures for all four obtained states, except a slight rotation of the cytoplasmic domains in [Na_3_]E2P. (B) Lipid density for wt and Q140L at a contour level of 3.5 (orange) and 4.5 (grey). (C) specific activity of wt and mutant Na,K-ATPases purified in the presence of dimerent lipid mixtures. (D) Inflection temperatures of wt and Q150L α1-Na,K-ATPase (equivalent of α3-Q140L) in the conformations shown on the x-axis. Additives in brackets were used to stabilise the conformation as shown in supp. Fig 1.

The mechanism of inactivation, especially the implication that the α3-Q140L mutation impairs the phospholipid binding in lipid site B, were elucidated from functional properties of α3-Q140L and the equivalent α1-Q150L (**Fig.3C and Fig. 4**). Na^+^,K^+^-ATPase activity of purified, detergent-solubilized, and lipid-bound complexes of α3, α3-Q140L, α1, and α1-Q150L, prepared with or without 18:0/22:6PE [62], showed the expected, approximately 1.5-fold stimulation of activity with 18:0/22:6 PE for wild type α3 and α1, but for α3-Q140L and α1-Q150L no such stimulatory effect was observed (**Fig. 3C**).

**Figure 4.**
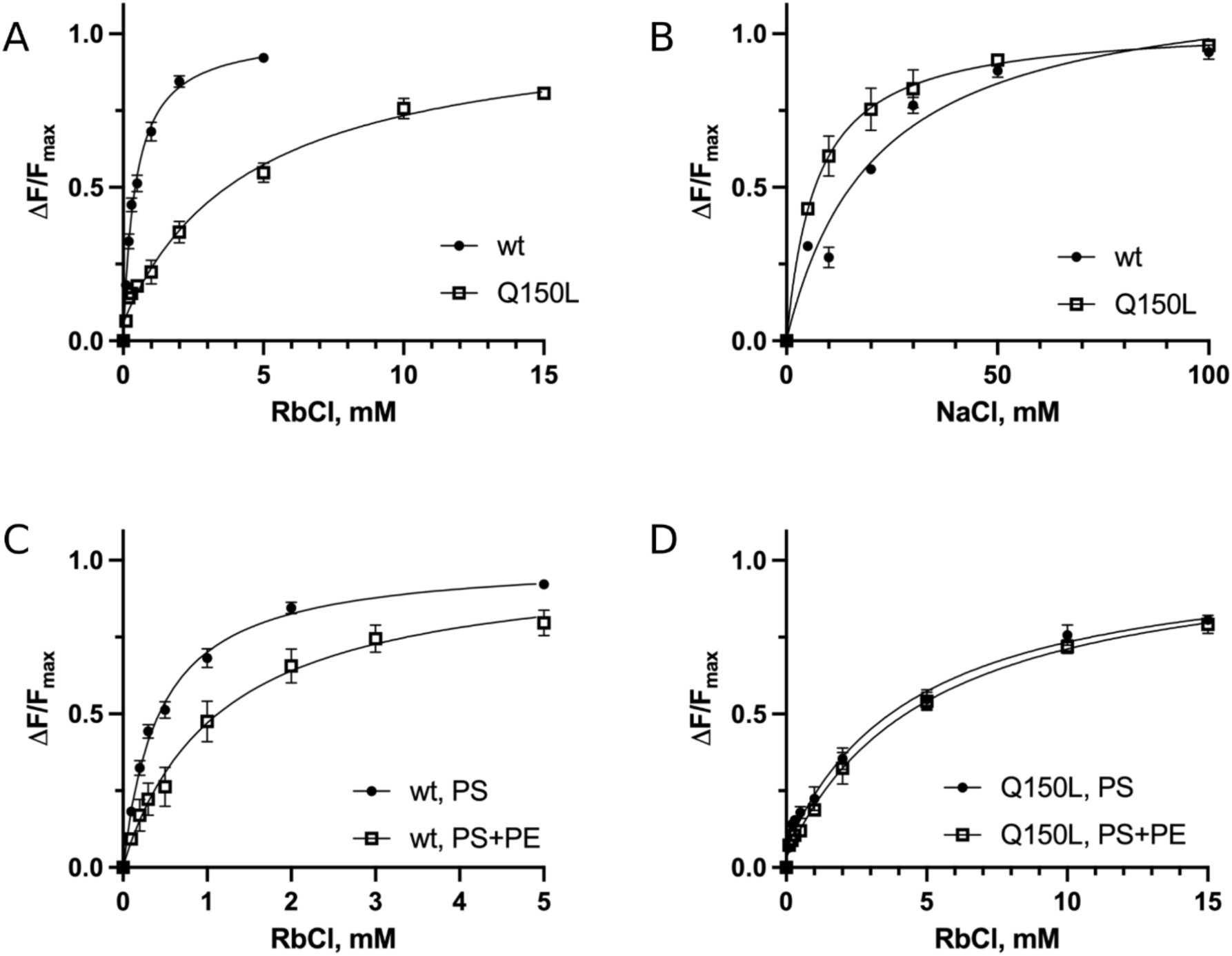
FITC-labelled Na,K-ATPase (α1β1). Cation titrations of fluorescence changes. The data represent averages ± SEM of 3-5 determinations at each Rb or Na concentration. Raised K_1/2_Rb with lower K_1/2_Na for Q150L versus WT, indicates relative poise of the E1/E2 conformational equilibrium towards E1. For α1WT, for PS+PE versus PS a similar conclusion pertains, but for the Q150L the similar K_1/2_Rb values for PS+PE versus PS suggest most simply that binding of PE is impaired. K_1/2_ values are as follows: Fig 4A & B WT: K_1/2_ Rb-0.43±0.02 mM; K_0.5_ Na− 16.0±1.0 mM, Q150L: Rb-2.78±0.27 mM; K_0.5_ Na− 6.6±2.3 mM, Fig 4C & D WT: PS-K_1/2_ Rb-0.43±0.02 mM; PS+PE-K_1/2_ Rb-1.12±0.48 mM, Q150L: K_1/2_ Rb-2.78±0.27 mM; PS+PE-K_1/2_ Rb-3.73±0.29 mM

Polyunsaturated PE is known to increase the rate of the E1→E2P transition [14]. Thus, if Q140L impairs the phospholipid-protein interaction, the overall E1:E2 balance should be affected. We examined this possibility using two complementary approaches. First, we compared thermal stability of WT α1β1FXYD1 and Q150L-α1β1FXYD1 in conditions known to stabilize different conformations, using nanoDSF (**Fig 3D, Suppl. Fig. 1**). Most conditions show similar inflection temperatures for WT versus mutant, but in conditions favouring E2P states, significantly lower inflection temperatures were observed for the mutant, suggesting that the E2P conformation was destabilized. Another approach used the FITC probe (see above) to assess the E1-E2 equilibrium of α1β1FXYD1 and Q150L-α1β1FXYD1, and the effect of 18:0/22:6 PE on K_0.5_(Rb) for transition to rubidium-bound E2 (**Fig. 4**). Indeed, cation titrations show that Q150L-α1 has a higher K_0.5_(Rb) for the E1 to E2 transition, and a lower K_0.5_(Na) for the reverse E2 to E1 transition, indicative of E2 conformations being favoured for WT and destabilised for the Q150L-α1 mutant (**Figs. 4A and B**). Finally, we compared WT and Q150L prepared with PS alone or with PS+18:0/22:6 PE. As seen in **Fig. 4C and 4D** PE significantly raises K_0.5_Rb for WT but has little or no effect on Q150L-α1. These findings all point consistently to the α1-Q150L and the equivalent α3-Q140L AHC mutation being shifted towards E1 conformations, due to an impaired interaction with the polyunsaturated PE at lipid site B.

Previous evidence for the specific functional roles of phospholipids to stabilize, stimulate or inhibit Na^+^,K^+^-ATPase in so-called sites A, B, and C respectively, has relied on observations *in vitro* using purified protein [13]. The evidence that the AHC-causing Q140L mutant is associated with impairment of specific 18:0/22:6 PE binding in lipid site B, provides convincing support for a role of this interaction *in vivo*. In addition, α3-Q140 is of interest as the residue is highly conserved among P-type ATPases (**Suppl. Fig. 7**). Indeed, an equivalent mutation has been reported for SERCA (Q108H) as causing Darier disease and reducing Ca^2+^-ATPase activity by ∼80% [64]. Also for SERCA, Q108 lines a phospholipid binding site [65]. An implication could be that phospholipid binding in this pocket regulates activity of several P-type ATPases. In native membranes, the proportion of polyunsaturated fatty acids in phospholipids (especially DHA) is positively correlated with molar activity of Na^+^,K^+^-ATPase, and is known to be an important factor in determining basal metabolic rate (reviewed [66]). The mechanism of this effect is unclear, but could reflect either general (e.g. via “membrane fluidity”) or specific lipid-Na^+^,K^+^-ATPase interactions. The observation that a point mutation in a specific site B impairs 18:0,22:6 PE binding and reduces Na^+^,K^+^-ATPase activity by about 80%, suggests that specific interactions of polyunsaturated fatty acid (PUFA) containing phospholipids are likely to play an important role in the physiological effects of the fatty acid unsaturation index [66].

## Conclusions

The cryo-EM structures and functional studies presented here depict the mechanisms of sequential partial reactions of the Na^+^ transport cycle **(Fig 5)**, favoured in forward reactions by a high ATP-to-ADP ratio. Physiologically, this proceeds until electrochemical gradients reach maximum [67], or regulation blocks function. Energy from ATP-hydrolysis is not transferred in a single stroke but over the whole cycle [68], and following Na^+^ dependent phosphorylation and ADP release, the newly observed [Na_3_]E2P conformation appears as the intermediate for which Na^+^ release is initiated in the E1P-E2P transition, orchestrating the changes in the tilt and displacement of the trans-membrane segments and breaking the cooperativity of Na+ sites that allow Na^+^ ions to dissociate at the extracellular side. The close approach of the P and A domains in [Na_3_]E2P, characteristic of E2 conformations, is the result of Na+ dependent phosphorylation and ADP release and may be triggered initially by charge neutralization of Asp366 in [Na_3_]E1P (including also bound Mg^2+^), which reduces electrostatic repulsion with Glu211 [69]. This creates strain in the linker regions that connect the cytoplasmic domains to the transmembrane domain that subsequently is released by rearrangement of the TM1-6 segment.

**Figure 5.**
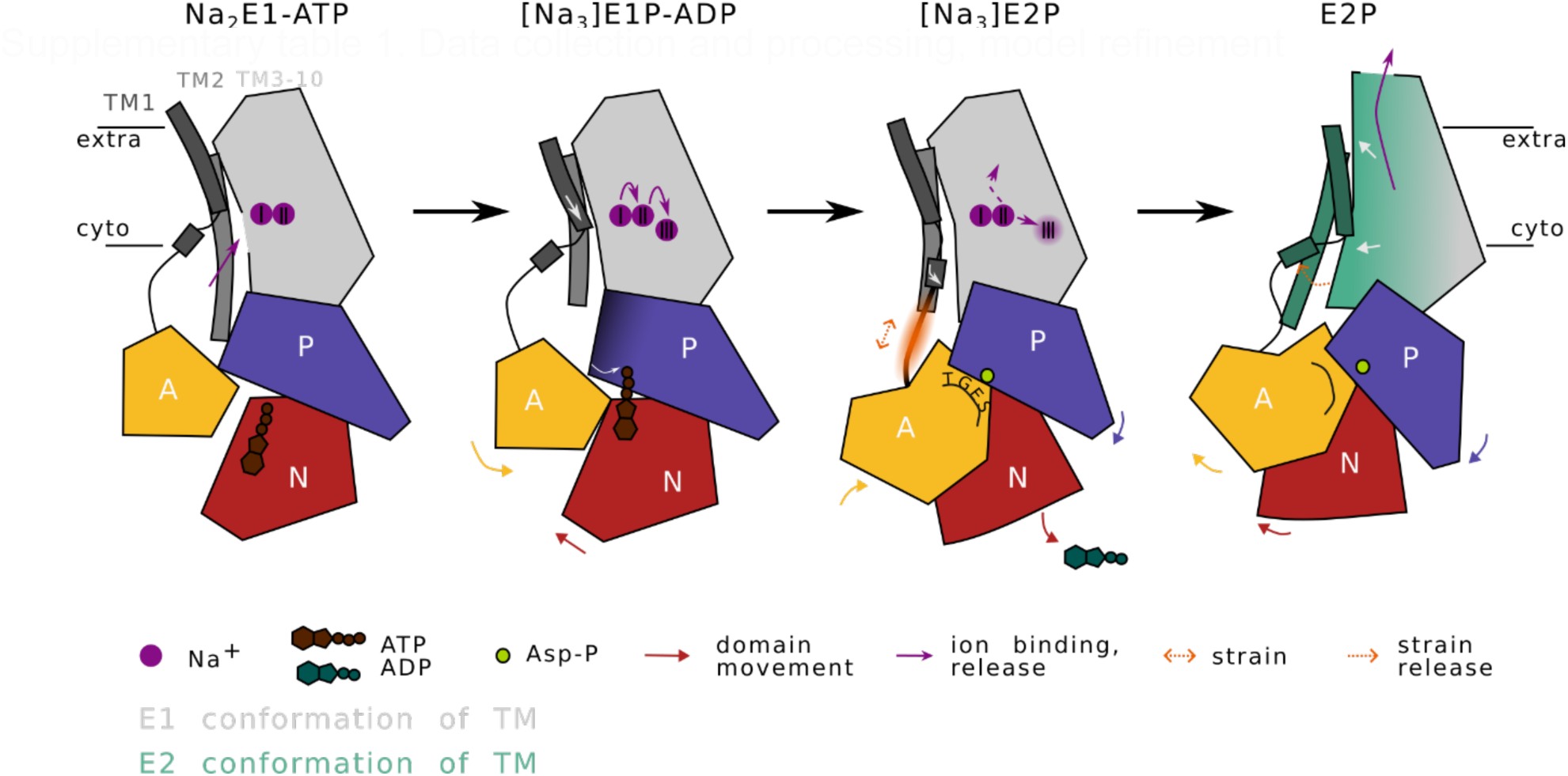
Schematic representation of principal rearrangements between structures obtained in the presence of Na and ATP. Arrows indicate changes to the previous conformation. Colouring as in previous figures. The three principal transitions are (1) binding of the 3^rd^ Na− ion pushes the two bound Na− ions to their neighbouring sites causing closure of the cytoplasmic gate and ion occlusion and causes the cytoplasmic domains to form a tighter arrangement allowing phosphorylation from ATP, (2) The E1P->E2P conformational transition causes an immediate rearrangement of the A and N-domain with the N-domain rotating backwards allowing the release of ATP and the A-domain rotation to cover the aspartyl-phosphate with its signature TGES-loop. The transmembrane domain remains largely unaltered except for destabilisation of site III and the straightening of TM1. (3) Destabilisation of site III and release of strain causes the rearrangement of TM1-6 while A, N and P-domain rotate in a concerted action.

A further significant finding is how α3 is adapted for physiological regulation of cytoplasmic Na^+^. Specifically, the lower apparent Na^+^ affinity of α3 compared to α1 is the result of a difference in intrinsic conformational equilibrium between E1 and E2 conformations, with the α3 isoform being poised towards E2, thus amplifying K^+^/Na^+^ competition at the cytoplasmic side. α3 becomes activated primarily upon intense nerve action and the associated increase in cytoplasmic Na^+^ concentration, thus serving to moderate the increase during nerve action and restore the Na^+^ concentration towards resting levels.

The spectrum of *ATP1A3* associated disorders is broad and ranges from moderate motor impairment to severe infantile epilepsy [61, 70]. Mutations leading to severe phenotypes are predominantly found near the ion binding sites [11]. A lack of Na^+^,K^+^-ATPase activity is not always sufficient to explain the broad phenotypic spectrum [71], and alternative mechanisms such as temperature instability [72], misfolding [73] and gain of function leak currents [41] have been proposed. Here we point to an impaired mechanism of lipid-based activation for the Q140L mutation, which also becomes destabilized in E2 conformations, raising the question whether destabilization of E2 states is a more general mechanism in AHC. The concept of impaired protein-lipid interaction in AHC suggests a possible strategy for treatment, by modification of membrane phospholipid content/saturation, or with lipid-like molecules acting as corrector compounds.

## Acknowledgements

We are grateful for excellent laboratory assistance by Anna Marie Nielsen and Karen Bech-Pedersen, for technical assistance by Taner Drace, Andreas Bøggild and Thomas Boesen of the Danish national cryo-EM facility EMBION, and by Jesper Karlsen for resources and assistance with scientific computing and data storage at the EMCC data science facility at Aarhus University. We thank Drs. Ronald J. Clarke, Natalya Fedosova and Joseph A. Lyons for valuable discussion. This work was supported by an Horizon2020 Marie Sklodowska Curie Actions fellowship to MH (793086), a professorship grant from the Lundbeck Foundation to PN (R310-2018-3713), an infrastructure grant from the Danish Ministry for Research and Higher Education (EMBION - 5072-00025B) and the Novo Nordisk Foundation (NNF20OC0060483), equipment grants from the Carlsberg Foundation (CF22-1535 and CF23-1394), and the CryoNet network supported by the Novo Nordisk Foundation.

## Data availability

Coordinates for the structures have been deposited in the Protein Data Bank (PDB) with the following accession codes: 1) 9RON, 9ROO, 9ROP, and 9RPQ for Na_2_E1-ATP, [Na_3_]E1P-ADP, [Na_3_]E2P and E2P states, respectively, of α1β1-FXYD1 at Na^+^ turn-over conditions; 2) 9RO9, 9ROA, 9ROD, and 9ROE for Na_2_E1-ATP, [Na_3_]E1P-ADP, [Na_3_]E2P and E2P states, respectively, of α3β1-FXYD1 at Na^+^ turn-over conditions; 3) 9ROF, 9ROG, 9ROH, and 9ROI for Na_2_E1-ATP, [Na_3_]E1P-ADP, [Na_3_]E2P and [K_2_]E2.P states, respectively, of α3β1-FXYD1 at full turn-over conditions (Na^+^ and K^+^); and 9ROJ, 9ROK, 9ROL, and 9ROM for Na_2_E1-ATP, [Na_3_]E1P-ADP, [Na_3_]E2P and E2P states, respectively of α3-Q140Lβ1-FXYD1 at Na+ turn-over conditions. Maps will be deposited at the EMDB.

## Materials and methods

### Saposin A expression and purification

Saposin A was expressed in *E. coli* Rosetta gami-2 (DE3) cells, transformed with a pNIC-Bsa4 plasmid encoding saposin A with an N-terminal TEV cleavable His_6_-tag. Expression and purification were performed essentially as described in Frauenfeld et al. [18]. Cells were grown in terrific broth medium (2% w/v LB, 2% yeast extract, 0.8% v/v glycerol, 70mM KH_2_PO_4_ pH 7.2) supplemented with 25µg/ml Kanamycin, 17µg/ml Chloramphenicol and 6µg/ml Tetracyclin to OD_600_ 0.6. Expression was induced by addition of 0.5mM IPTG and continued overnight at 20°C prior to cell harvest by centrifugation. The cells were resuspended in 4ml buffer/g cells (20mM MOPS/Tris pH 7.4, 150mM NaCl, 5mM Imidazole, 1mM PMSF), disrupted by sonication and debris removed by centrifugation (20,000g 30min). Supernatant was heated to 85°C (10min) and subsequently clarified by centrifugation (20,000g 30min). Saposin A was purified by IMAC, incubating the supernatant with 1ml Ni Sepharose/10g cells (Cytiva) at RT for two hours. Resin was transferred to a gravity flow column and washed with 10x bead volume wash buffer (20mM MOPS/Tris pH 7.4, 150mM NaCl, 20mM Imidazole). Saposin A was eluted in wash buffer supplemented with 200mM imidazole. TEV protease and 1mM DTT (saposin A:TEV, 1:100(w/w) was added and the sample dialyzed overnight against 20mM MOPS/Tris pH 7.4, 150mM NaCl. TEV protease was removed by reverse IMAC and Saposin A was purified further by size exclusion chromatography (column: Superdex 200 pg, 16/600, buffer: 20mM MOPS/Tris pH 7.4, 150mM NaCl) and fractions containing monomeric protein were pooled, flash frozen in liquid nitrogen and stored for at −80°C.

### FXYD1 expression and purification

The regulatory FXYD1 subunit, with a N-terminal His_6_-tag, was expressed in *E. coli* C41(DE3) cells and purified effectively as described in [74]. Cells were grown in LB medium to OD_600_ 0.8 and expression induced with 0.5mM IPTG. Cells were cultured over night at 18°C and harvested by centrifugation. The cell pellet was resuspended in 5 ml buffer/g cells (20mM MOPS/Tris pH 7.4, 500mM NaCl, 5mM MgCl_2_, 1mM EDTA and 1mM PMSF) and disrupted by sonication. Membranes and inclusion bodies were isolated by centrifugation (200,000g, 30min). The pellet was solubilized in 15ml/g solubilization buffer (10mM MOPS/Tris pH 7.4, 500mM NaCl, 10%Glycerol, 5mM Imidazole, 4mg/ml DDM, supplemented with 8M Urea) for one hour at RT, before dilution to 6M Urea and debris removal by centrifugation (200,000g, 30min). The supernatant was incubated for one hour with 2.5ml of Ni Sepharose/10g cells, transferred to a gravity flow column and the resin washed with 10x bead volume wash buffer (10mM MOPS/Tris pH 7.4, 500mM NaCl, 10%Glycerol, 5mM Imidazole, 1mg/ml DDM, 4M Urea). FXYD1 was eluted in wash buffer with 0.3mg/ml DDM supplemented with 200mM imidazole and dialyzed overnight against 10mM MOPS/Tris pH 7.4, 500mM NaCl, 10% Glycerol, 0.1mM DTT in the presence of TEV protease. The protease was subsequently removed by reverse IMAC and purified FXYD1 collected and flash-frozen in liquid nitrogen.

### Na^+^,K^+^-ATPase (α_1_β_1_) expression, purification and saposinA nanodisc reconstitution

Human Na^+^,K^+^-ATPase isoforms and mutants were expressed in *Pichia pastoris/Komagataella phaffii* SMD1165 (*his4, prb1*) yeast by methods described previously with some modifications as follows [14, 75, 76]. Cells were grown in BMG medium (100mM KP_i_ pH 6, 3.4g/L YNB (wo. Amino acids), 10g/L (NH_4_)_2_SO_4_, 0.4mg/L Biotin, 0.4% glycerol (v/v)) to OD_600_ 9 in a bench-top bioreactor BIOFLOIII (New Brunswick Scientific) at 30°C. A fed-batch feeding procedure from a 20% glycerol stock was initiated till OD_600_ −25.Temperature was reduced to 25°C and expression induced by repeated supplementation of 1% MeOH with 12 hours interval. Expression was continued for 5 hours after the second induction before cell harvest by centrifugation (1,000g, 10min). The cell pellet was suspended in lysis buffer (40mM MOPS/Tris pH 7.4, 1mM EDTA, 1.2M Sorbitol) and lysed by mechanical shear utilizing a Bead-Beater. Debris and unbroken cells were removed by centrifugation (5,000g, 20min) and the membrane containing fraction collected by ultracentrifugation (180,000g, 90min) of the supernatant. The membrane containing pellet was suspended in 1ml buffer/g cells (20mM MOPS/Tris pH-7.4, 1mM PMSF, 2M Urea), incubated for 20 min and the membranes collected by a second round of ultracentrifugation. Supernatant was discarded, the pellet suspended to 300mg/ml w/v in suspension buffer (20mM MOPS/Tris pH −7.4, 20% glycerol, 1mM PMSF and 5ug/ml Chymostatin, Pepstatin and Leupeptin), flash-frozen in liquid nitrogen and stored at −80°C.

Purification was initiated by dilution of the membrane to 100mg/ml in solubilization buffer (20mM MOPS/Tris pH 7.4, 200mM NaCl, 10% glycerol, 20mM Imidazole) with subsequent addition of DDM/cholesterol hemisuccinate (10/1% w/v) to a membrane:DDM ratio of approx. 11:1 (w/w). The solution was sonicated for 1 min and unsolubilized material removed by centrifugation (180,000g, 30min). The supernatant was incubated with TALON resin (1ml resin /4-5g total membrane) for 5 hours, 90% of the supernatant removed and 55μg FXYD1 / g membrane was added and incubated overnight. Unbound material was removed and the resin washed with 10x bead-volume solubilization buffer supplemented w. 0.5mg/ml C12E8 before suspending the resin in an equal volume of relipidation buffer (solubilization buffer + 0.5mg/ml C12E8, 0.15mg/ml SoyPC, 0.15mg-ml SOPS, 0.05mg/ml Cholesterol, 3mM MgCl_2_ and 1mM ATP). Nanodisc reconstitution of the resin bound Na^+^,K^+^-ATPase was performed by addition of 1.35ml 1mg/ml saposin A per. ml resin, 20 min incubation and subsequent wash with 3x bead-volume solubilization buffer without detergent. The protein was eluted in solubilization buffer supplemented with 200mM imidazole, concentrated and further purified by size exclusion chromatography utilizing a Superdex 200(10/300)-increase column equilibrated in 20mM MOPS/Tris pH 7.4, 200mM NaCl, 3mM MgCl_2_.

### Sample preparation and EM data acquisition

Purified monomeric nanodisc reconstituted Na^+^,K^+^-ATPase was diluted to approx. 0.5mg/ml in SEC buffer supplemented with ATP and LMNG to final concentrations of 1mM and 1.5CMC, respectively. 3µl sample were immediately put on freshly glow-discharged c-Flat 1.2/1.3-300 copper grids (Protochip) (45sec at 15mA using a PELCO easiGlowTM), blotted on a Vitrobot Mark IV (ThermoFisher Scientiffic) with a 10sec wait time, 5.5sec blot time at blot force 1 and plunge-frozen in liquid ethane.

For α1β1 (wt) data was collected at eBIC (electron Bioimaging Centre, Dimond) on a Titan Krios equipped with a K3 detector (Gatan) operated at 300kV. ThermoFisher Scientific EPU was used for automated data acquisition operating with a pixel size of 0.41Å and total exposure dose of approx. 60 e-/Å2 (**Suppl. Table 1**). The wt α3β1-FXYD1 data sets were collected at the Danish National Cryo-EM Facility (EMBION) on a Titan Krios equipped with a K3 detector at 300KV and a nominal dose of 60e/Å2 at a pixel size of 0.66Å. All datasets were collected using aberration free image shift.

### Cryo-EM image processing

The dataset was imported to and processed in cryoSPARC v4 [77]. For detailed processing workflow see **Supplementary Figure 8**. In short, motion correction and CTF estimation was performed and a subset of the data pre-process by blob picking and 2D classification. The best 2D class averages representing the reconstituted Na^+^,K^+^-ATPase in multiple orientations were used for reference based template picking, with subsequent 2D class averaging. Using ab-initio models generated from a subset of particles, the dataset was iteratively refined by hetero-refinement against 3-4 ab-initio models. The reduced particle stack was further refined by 3D classification without mask. Particles from volumes showing well resolved features were re-extracted and further processed by non-uniform refinement and 3D classification with a focussed and solvent mask. Selected volumes were re-run through non-uniform refinement and resolution estimated by GSFSC at 0.143.

### Model building and validation

Model building of the structures was initiated by docking the best fitting published crystal structures of native Na^+^,K^+^-ATPase (pdb 3WGU and 4HYT) into the obtained maps. Using SWISS-model homology modelling [78] the sequence of the models was converted into the human α_3_β_1_-FXYD1 or α_1_β_1_-FXYD1. The final models were generated through fitting in Namdinator [80] and iterative rounds of manual building in Coot [79] and real space refinement in Phenix [81]. Figures were prepared using ChimeraX [82].

### ATPase activity measurement

ATPase activity was measured using the Baginsky method [83, 84], measuring the amount of free inorganic phosphate generated by turnover of the Na^+^,K^+^-ATPase. The nanodisc reconstituted protein was incubated at 37°C in a preheated reaction mixture (20mM MOPS/Tris pH 7.4, 120mM NaCl, 20mM KCl, 1 mM ATP, 2mM MgCl_2_, 1mM EGTA) and 50μl aliquots transferred at selected timepoints to an equal volume of stop/development solution: (5:1 (v/v) mixture of solution A: 0.17M ascorbic acid, 0.1% SDS dissolved in 0.5M HCL and solution B: 28.3mM ammonium heptamolybdate). After a subsequent 10min incubation, 70μl 68 mM trisodium citrate, 154 mM sodium (meta)-arsenite, 2%(v/v) acetic acid were added, and the solution incubated for 1hour [85]. Absorbance at 860nm was measured using a Wallac Victor 3 Multilabel counter (Perkin Elmer). For each assay a standard curve of known Pi concentrations was included and activity calculated from the linear slope.

### Differential scanning fluorimetry

Thermostability was measured by nanoDSF using a Nanotemper Prometheus Panta operating at a temperature gradient from 20-95°C at 1°C/min. Samples were purified as described above in buffer containing choline chloride instead of NaCl. For stabilisation of selected conformations, the following 5x mixtures were prepared (all with 15 mM MgCl_2_): (1) sodium-bound E1 - 1M NaCl, (2) sodium-bound E1-ATP - 1M NaCl, 5mM AMPPCP, (3) [Na_3_]E1P-ADP - 1M NaCl, 25mM NaF, 5mM AlCl_3_, 5mM ADP, (4) E2P - 25mM NaF, 5mM BeSO_4_, (5) [K_2_]E2P - 1M KCl, 25mM KF, 5mM AlCl_3_, (6) [K_2_]E2.P_i_ - 1M KCl, 25mM KF, 5mM MgCl_2_, (7) [K_2_]E2.P_i_-ATP - 1M KCl, 25mM KF, 5mM MgCl_2_, 5mM AMPPCP, (8) [K_2_]E2-ATP - 1M KCl, 5mM AMPPCP, (9) [K_2_]E2 - 1M KCl. Inflection points were determined from first derivative of the 350/330nm ratio.

### FITC labelling and equilibrium titrations

FITC labeling of recombinant Na^+^,K^+^-ATPase in *P. pastoris* plasma membrane was performed essentially according to [69]. Briefly, membranes were first incubated with 0.3mg/ml SDS for 20’ at 24°C in a medium containing: 20 mM MOPS-Tris pH 7.4 + 10% glycerol to remove extrinsic proteins. Membranes were washed and pelleted, resuspended at 2mg/ml, incubated with 1 μM FITC in a medium containing: 50 mM NaCl, 1 mM EDTA and 20 mM TAPS-Tris pH 9.2, for 1 h, at 0°C. The reaction was stopped by addition of 6 volumes of cold: 100 mM MOPS-Tris pH 6.45 for 10’ on ice, followed by centrifugation at 150,000 g for 40 min. The pelleted labeled membranes were resuspended in 10 mM MOPS-Tris, pH 7.4, and 10% glycerol. The labeled membranes were solubilized and Na^+^,K^+^-ATPase protein was purified as previously described [47].

Fluorescence measurements were carried out in a Cary Eclipse spectrophotometer (Varian, Victoria, Australia). Samples containing 5 ug protein were placed in 1 ml cuvettes in 0.8 ml medium containing: 150 mM choline-Cl, 10 mM HEPES-Tris, pH 7.5, 5 mM MgCl_2_, 0.5 mM EDTA. The sample was stirred continuously with a magnetic stirrer at a temperature of 24°C. Excitation and emission wavelength were set at 485 nm and 520 nm, respectively. Fluorescence titrations were performed by adding repeated 5µl aliquots of 1M RbCl until the fluorescence was stable. Na^+^ titrations were performed by first adding 20mM RbCl and then repeated 5µl aliquots of 2M NaCl. Conformational equilibria were determined by first adding 20mM RbCl until fluorescence was stable and then 100mM NaCl. Values of fluorescence deflections were corrected for dilution effects and the accumulated values were then plotted as functions of total Rb^+^ or Na^+^ concentrations. The curves were fitted to simple saturation functions to obtain K_0.5_ values. Each curve was repeated 3 to 5 times.

#### Plasmids construction for expression of mutations Q150L and Q150H in human α1 and Q140L in human α3

Mutations were introduced into α1 and α3 by inverse PCR, followed by ligation using the KLD kit (M0554S, NEB). The resulting mutated pHil-D2 vectors containing the following Na^+^,K^+^-ATPase variants: α1(Q150L)β1-FXYD, α1(Q150H)β1-FXYD and α3(Q140L)β1-FXYD. Correct integration and sequence was confirmed by sequencing. The pHIL-D2 vectors, containing the mutated genes, were linearized by Not I digestion and used to transform spheroplasts of *P. pastoris* SMD1165, and His^+^ Mut^s^ transformants were selected as described previously [76]. Membrane preparations were made, and clones were screened for optimal protein expression by Western blotting using the anti-KETYY antibody.

### Expression and purification of Na^+^,K^+^-ATPase isoforms and mutants

Large scale growth of isoforms was done in Bellco Spinner FlasksTM in 10-liter volumes of growth medium, expression of the Na^+^,K^+^-ATPase was induced by adding 0.5% methanol daily for 4 days 25°C. Cells were collected, washed, broken with glass beads, and membranes were prepared as described previously [76].

Solubilization of membranes in DDM followed by purification of the Na^+^,K^+^-ATPase enzymes have been described in detail in Katz et al [76]. All isoform complexes were eluted from BD Talon beads in a solution containing 250mM imidazole, 100mM NaCl, 20 mM Tricine-HCl, pH 7.4, 0.1 mg/ml C_12_E_8_, 0.08 mg/ml SOPS, 0.01 mg/ml cholesterol, and 25% glycerol. The proteins were stored in aliquots at −80 °C.

### Na^+^,K^+^-ATPase activity of purified isoform complexes and mutants

Purified enzymes were diluted to 0.1ug/100ul in standard reaction medium consisting of 130 mM NaCl, 20 mM KCl, 3 mM MgCl2, 25 mM histidine, pH 7.4, 1 mM EGTA, 1 mM ATP with added lipids (0.005 mg/ml C_12_E_8_, 0.01 mg/ml SOPS, 0.001 mg/ml cholesterol). Incubation was performed at 37^0^C for different times. Phosphate release was measured using a malachite green dye (Pi Color Lock, Innova Biosciences [76].

### Ouabain binding

[^3^H]Ouabain binding capacity of yeast membranes was assayed as described by Katz et al. [76]. An equivalent of 200 µg membrane protein was incubated for 1 hr at 37°C in 10 mM MOPS-Tris, pH 7.2, 3 mM MgCl2, 1 mM Tris-vanadate and 1mM EGTA supplemented with 400nM [^3^H]Ouabain (Perkin Elmer, 30–40 Ci/mmol). The reaction was stopped by dilution in cold buffer of Tris-HCl 10mM pH 7.4, and membranes captured by filtration on glass fiber filters. Unbound material was washed and the amount of bound [^3^H]Ouabain determined by scintillation counting.

## Supplementary Figures

**Supplementary Figure 1.**
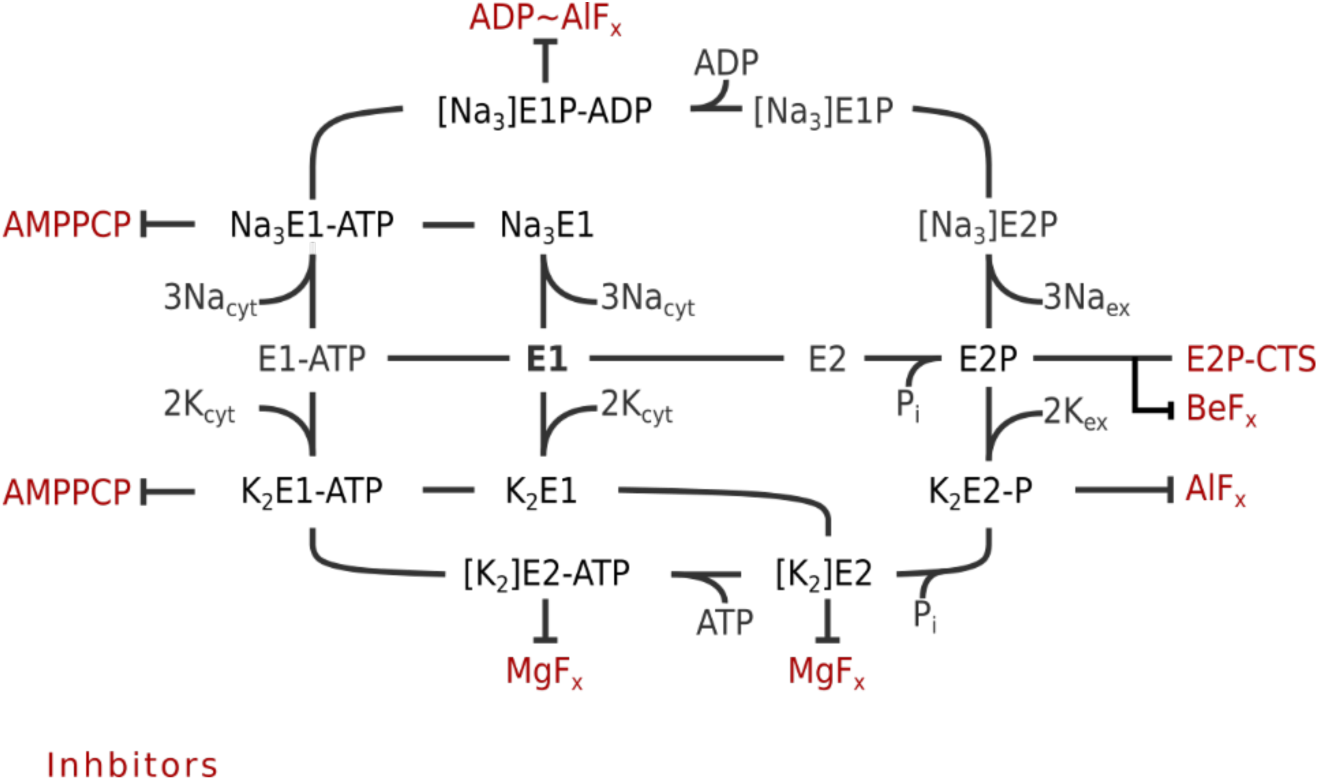
Post Albers cycle of Na,K-ATPase. Post Albers scheme of the Na,K-ATPase. Conformations with prefix [Na_3_] or [K_2_] indicate occluded conformations. Inhibitors shown in red can be used to stabilise respective conformations when applied with appropriate ions and have been used for nanoDSF analysis shown in figure 3D.

**Supplementary Figure 2.**
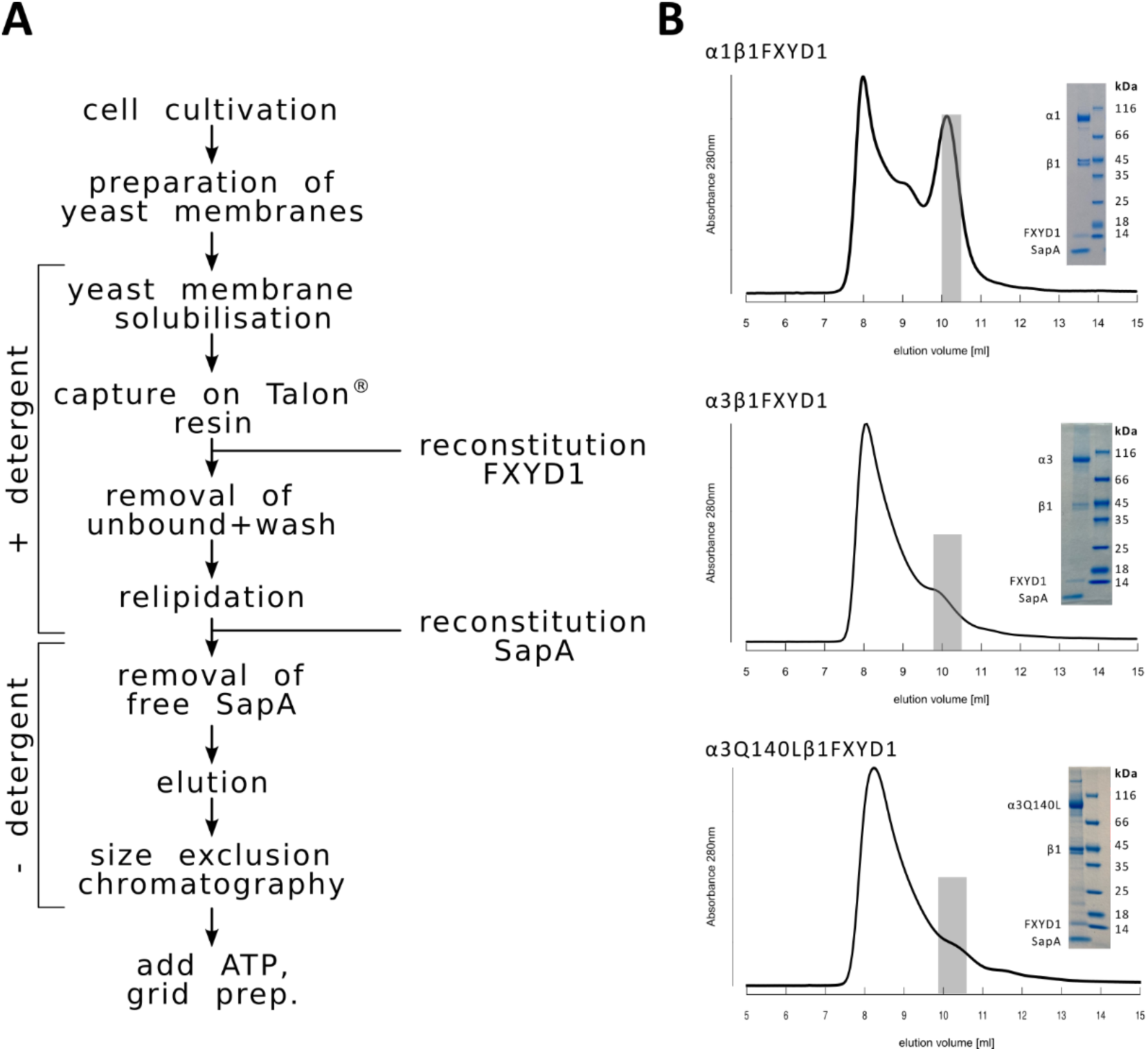
Purification of Na,KATPase in Saposin A nanodiscs. (A) Flowchart depicting the purification and reconstitution steps. (B) SDS-gels and size exclusion chromatograms of respective samples. Grey bars indicate fractions that were pooled and applied on grid.

**Supplementary Figure 3.**
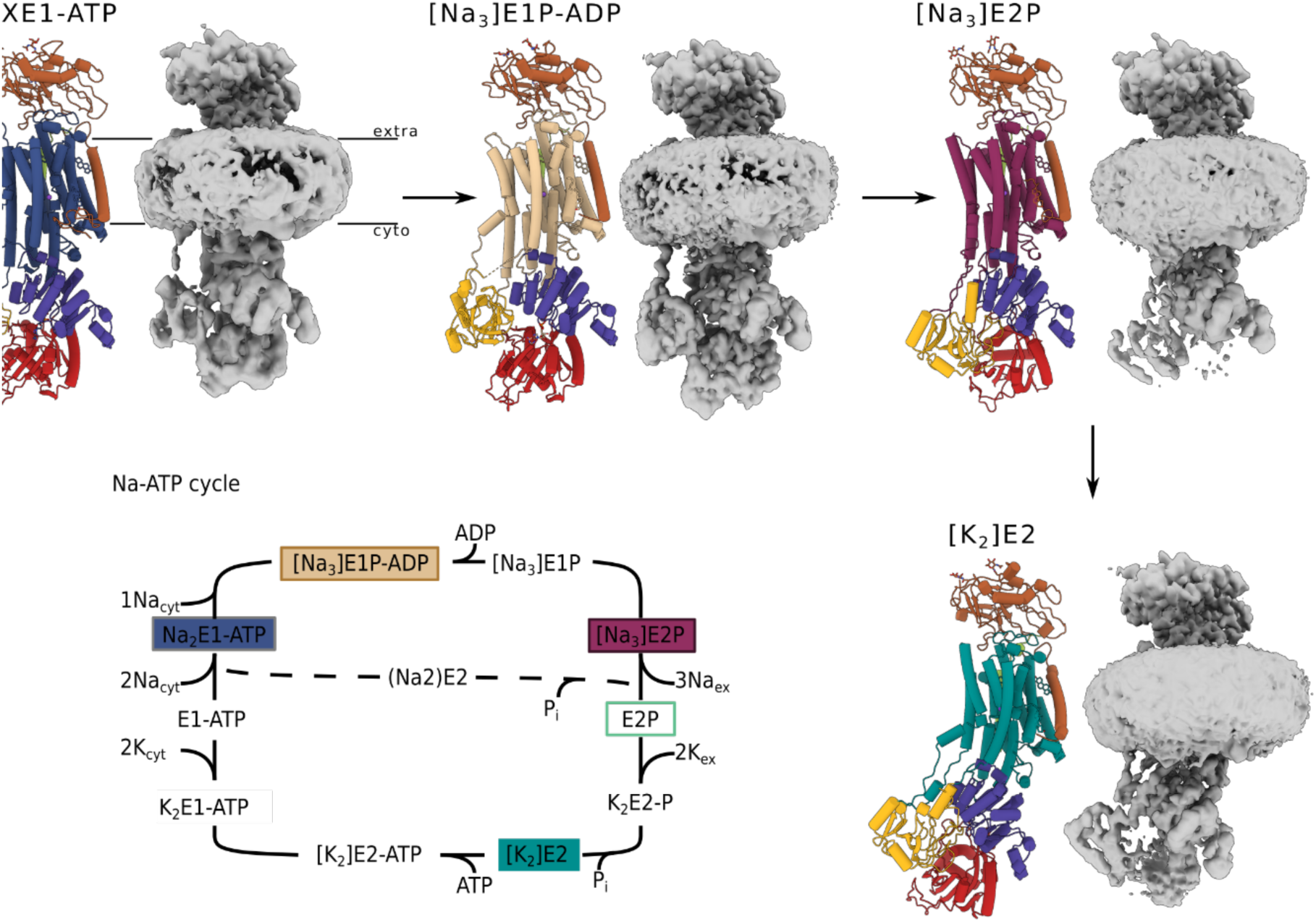
Cryo-EM of Na,K-ATPase under full turnover conditions in the presence of Na, K and ATP. Cryo-EM maps and structures of α3β1FXYD1 obtained in the presence of saturating concentrations of NaCl, KCl and ATP. Note that cations for the E1-ATP conformation have been donated X since it is unclear whether the state represents a Na^+^-, K^+^- or mixed Na^+^/K^+^-bound state. Under the given experimental conditions, a mix of all three is likely for this dynamic state. The Post-Albers scheme shows the conformation obtained under full turnover in filled coloured rectangles while conformations obtained under Na− only turnover are indicated in boxed rectangles.

**Supplementary Figure 4.**
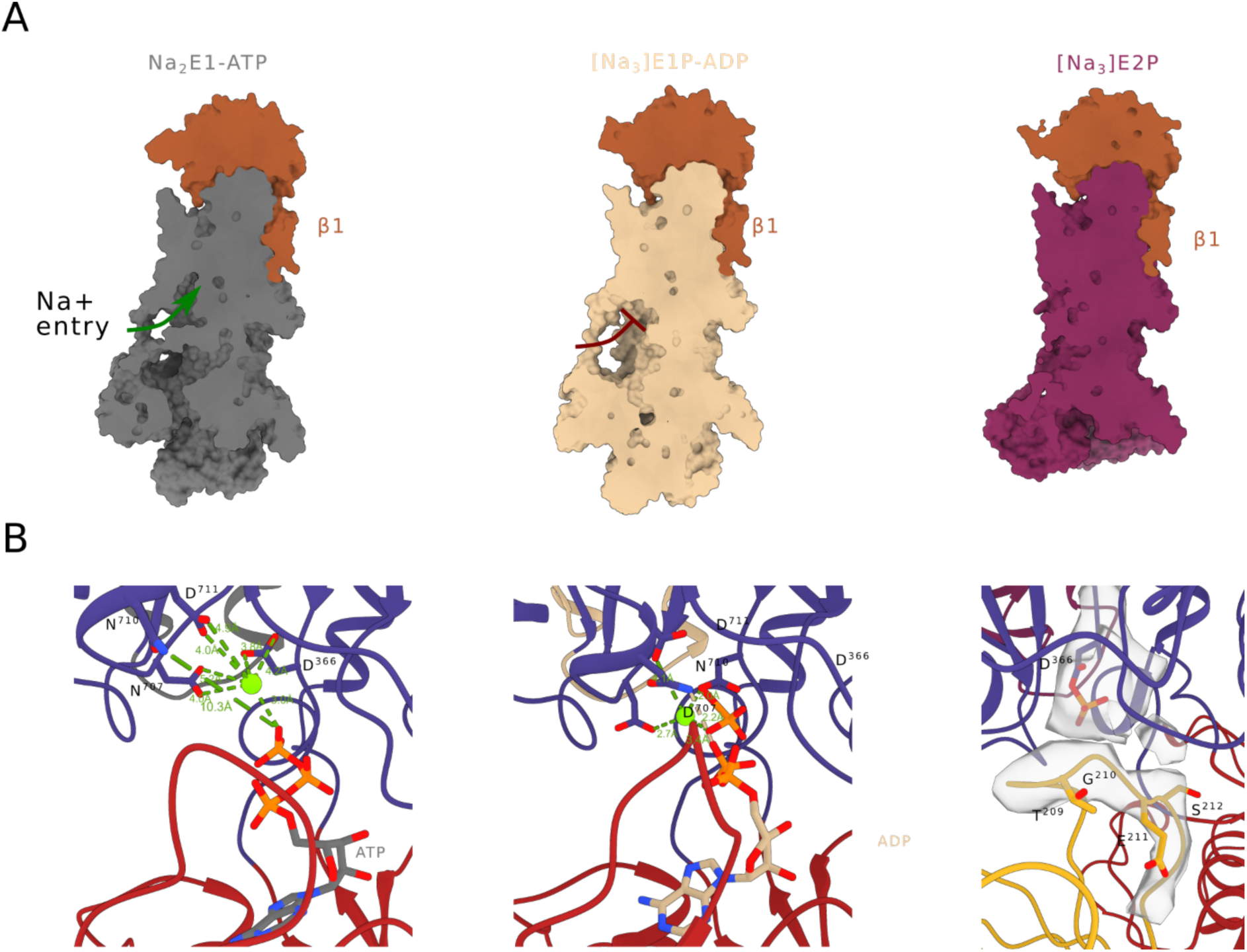
Comparison of Na^+^-bound conformations. (A) Clipped view of the Na^+^-bound states showing the opening towards the ion binding sites in E1 compared to the two occluded forms. (B) Close view on the phosphorylation site in the three Na− bound states.

**Supplementary Figure 5.**
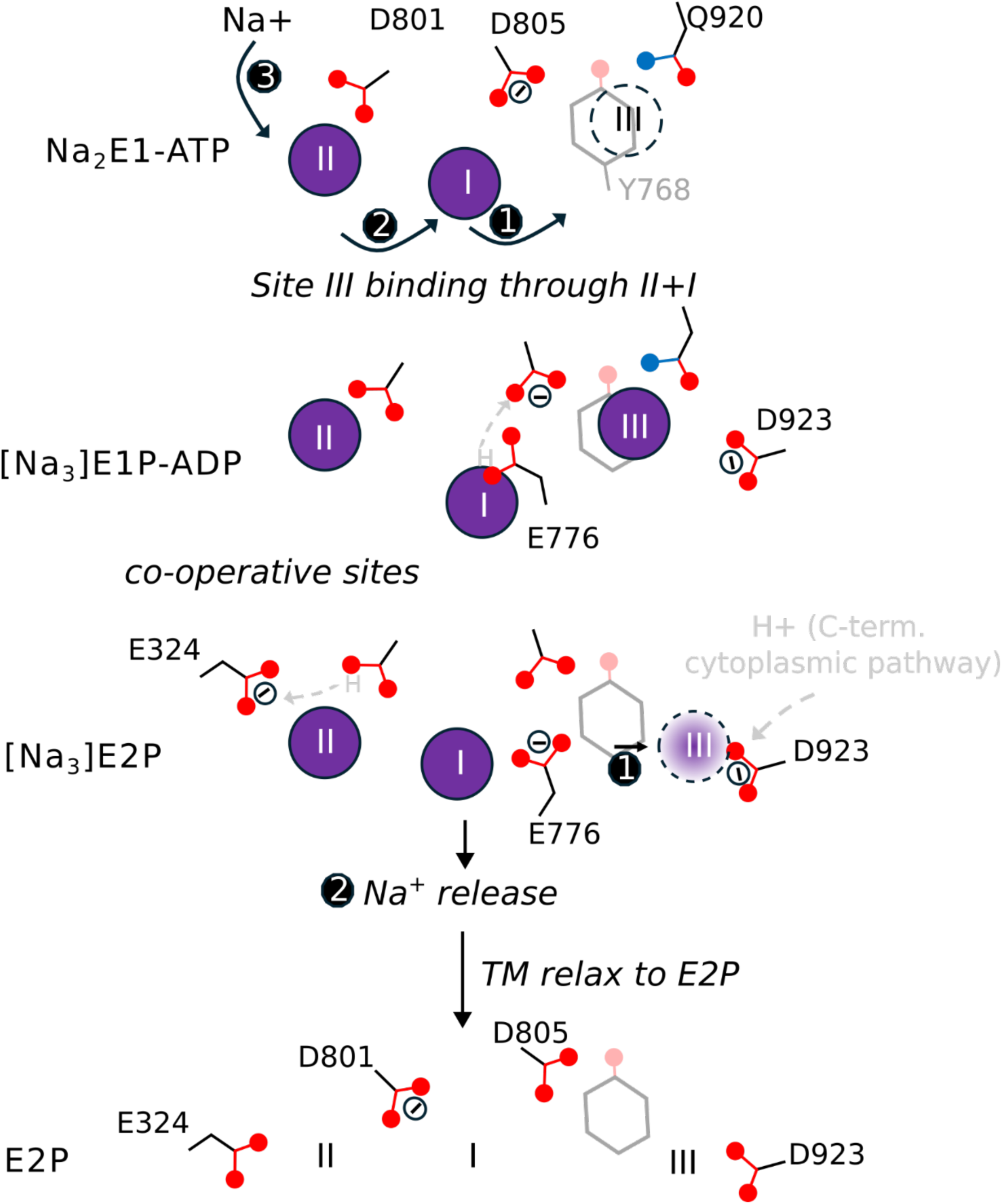
Sequence of Na-binding and release. Black arrows indicate movements of sodium ions (purple), grey arrows of protons. Deprotonated residues are indicated by Θ. The model is based on structural findings in this study with information on protonation of residues gathered from references 44-47.

**Supplementary Figure 6.**
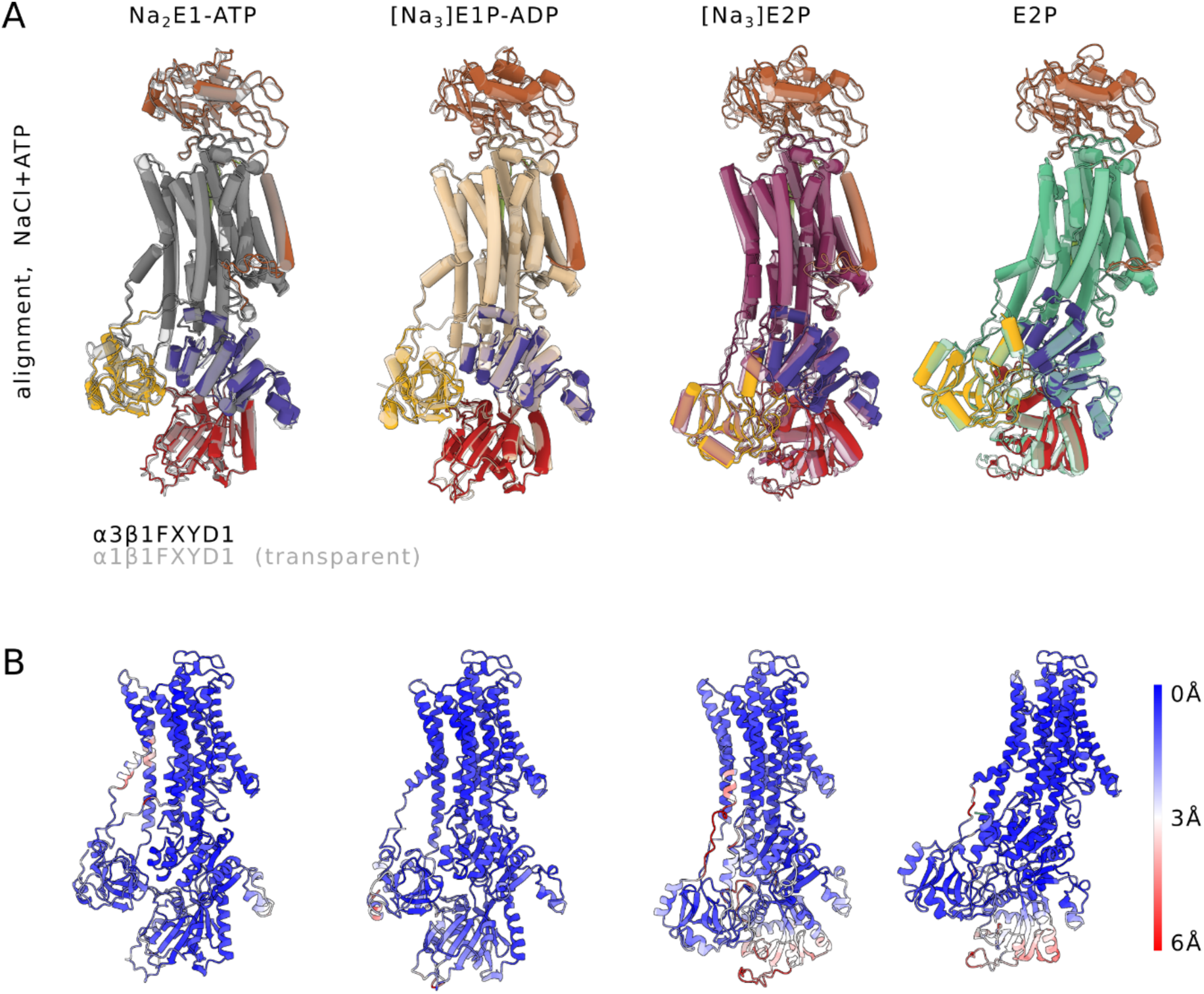
Comparison of Na,K-ATPase α1 and α3. (A) Alignment of α1 (semi-transparent) and α3 structures on the β-subunit. (B) Cα-RMSDs for alignments shown in (A).

**Supplementary Fig 7.**
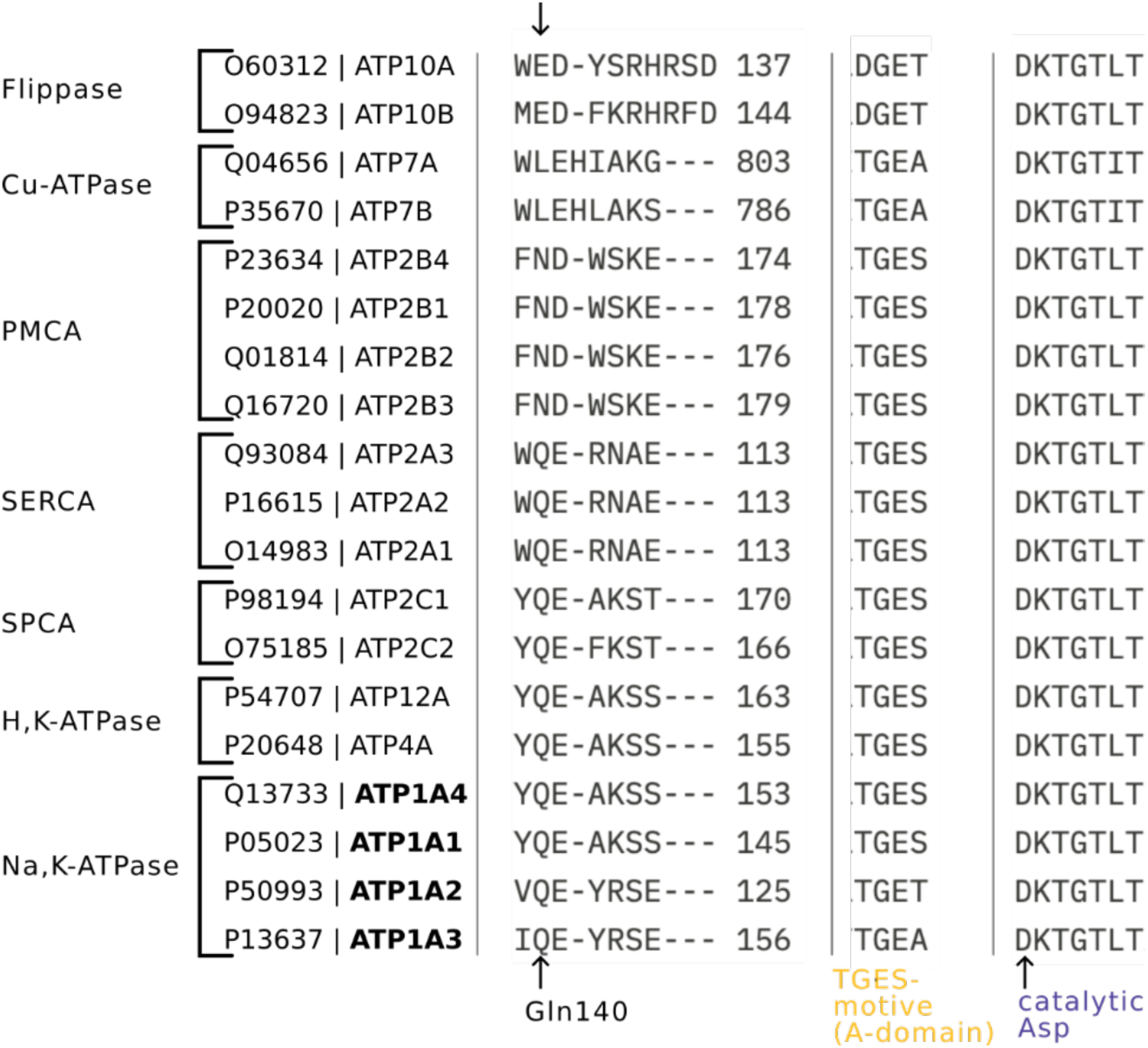
Sequence alignment of human P-type ATPases. The alignment was performed using the EMBL-EBI MSA clustal Omega site using indicated uniport IDs.

**Supplementary Fig 8.**
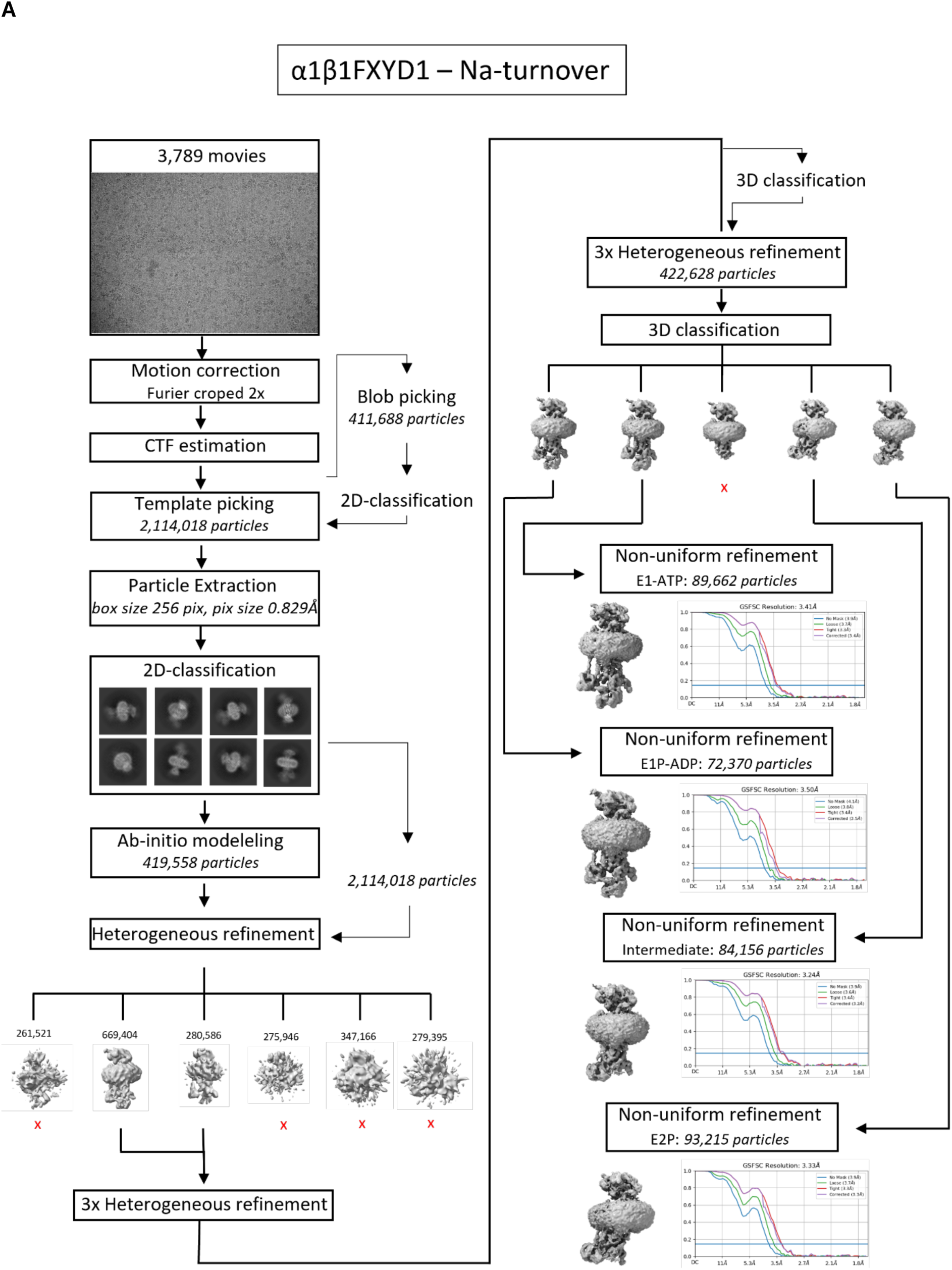
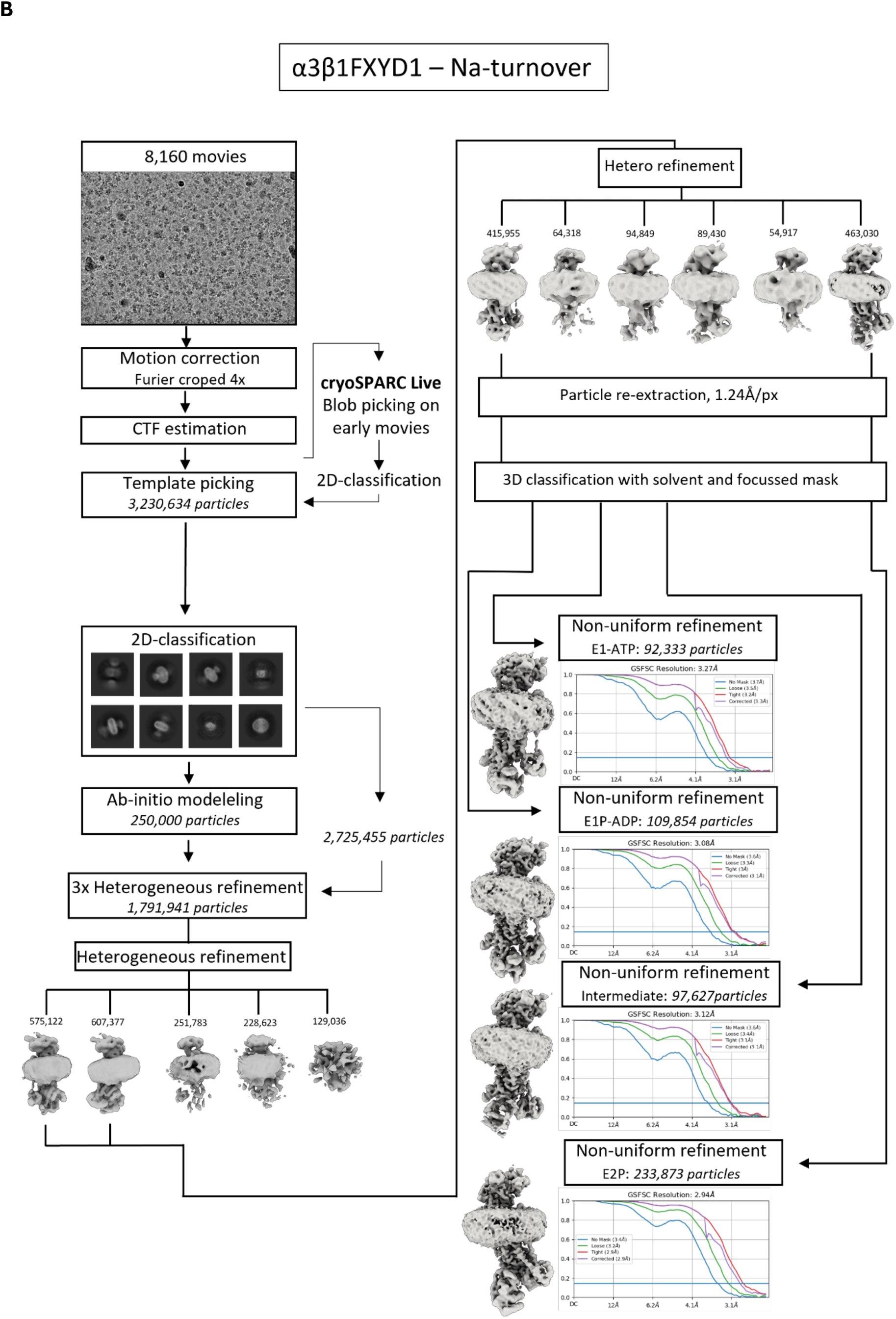
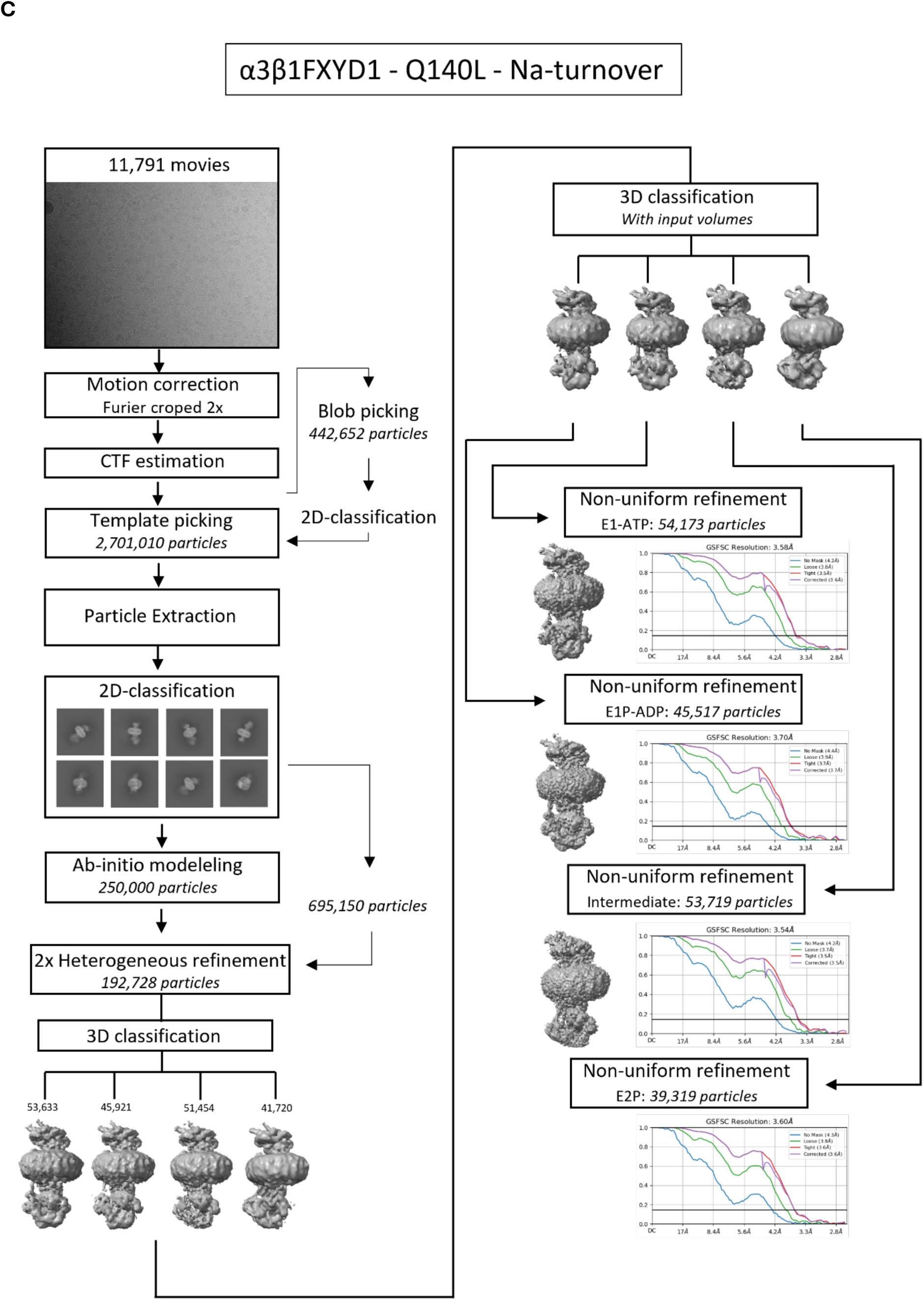
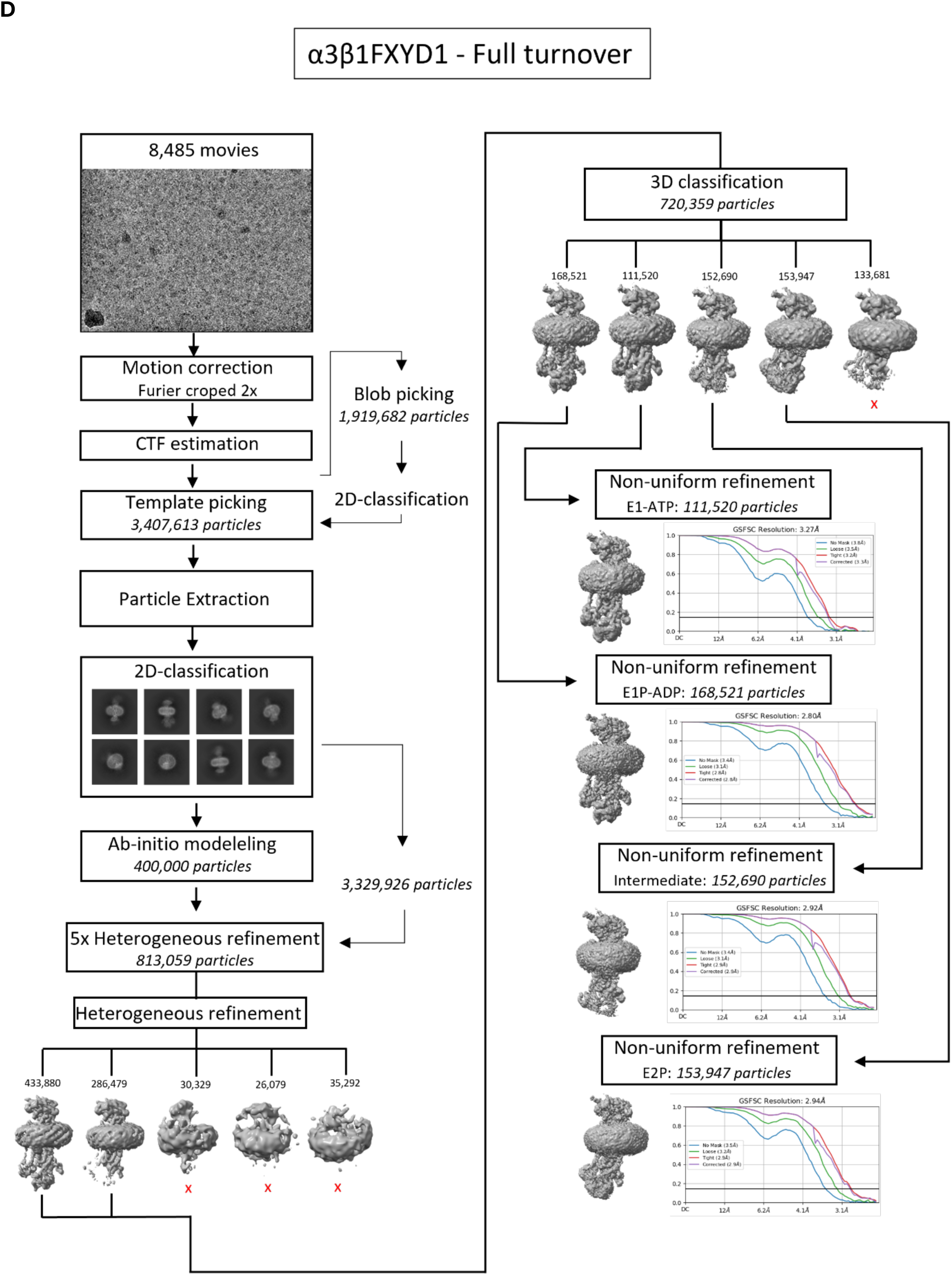
CryoSparc processing workflows.

**Supplementary 1.**
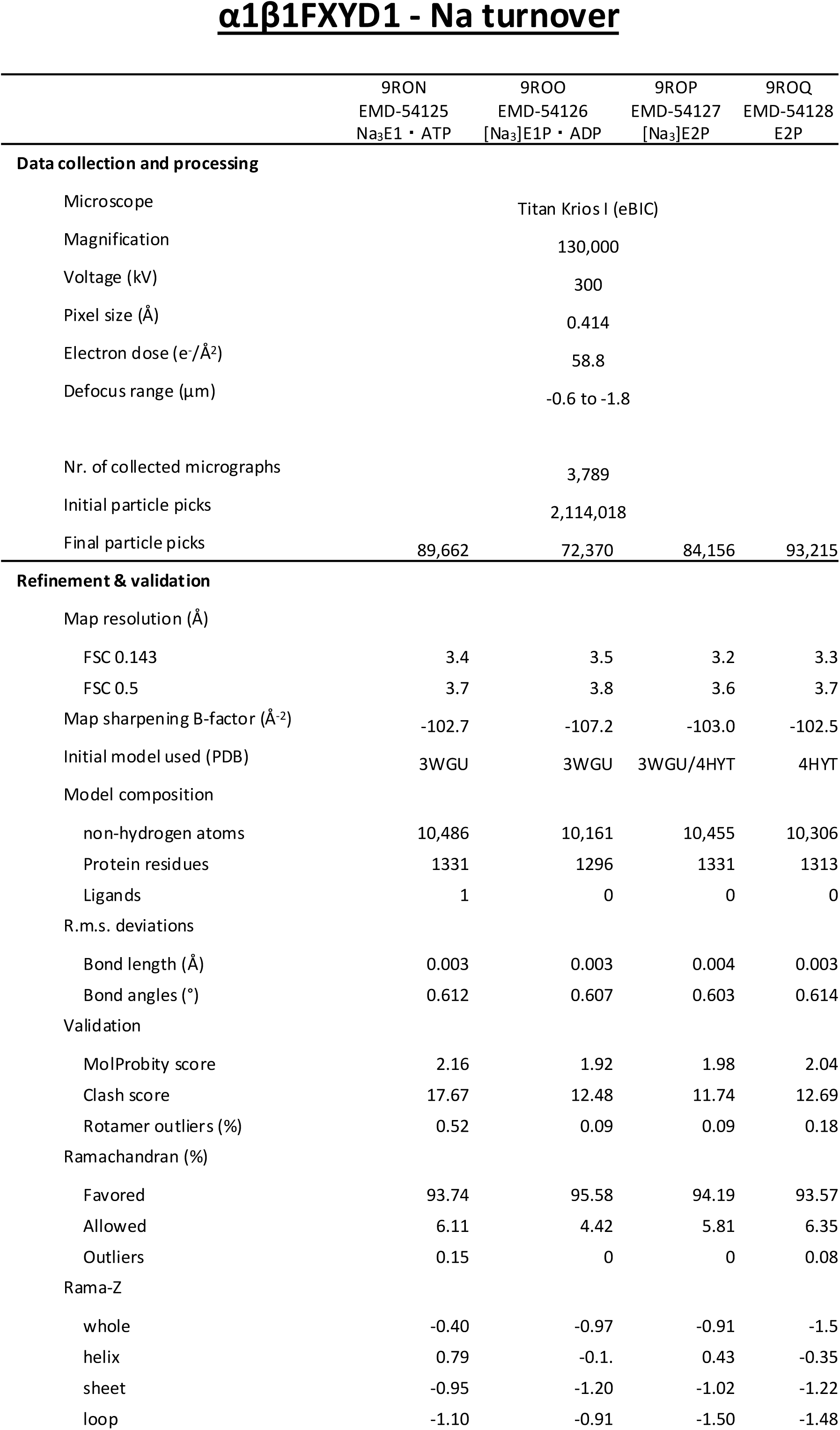
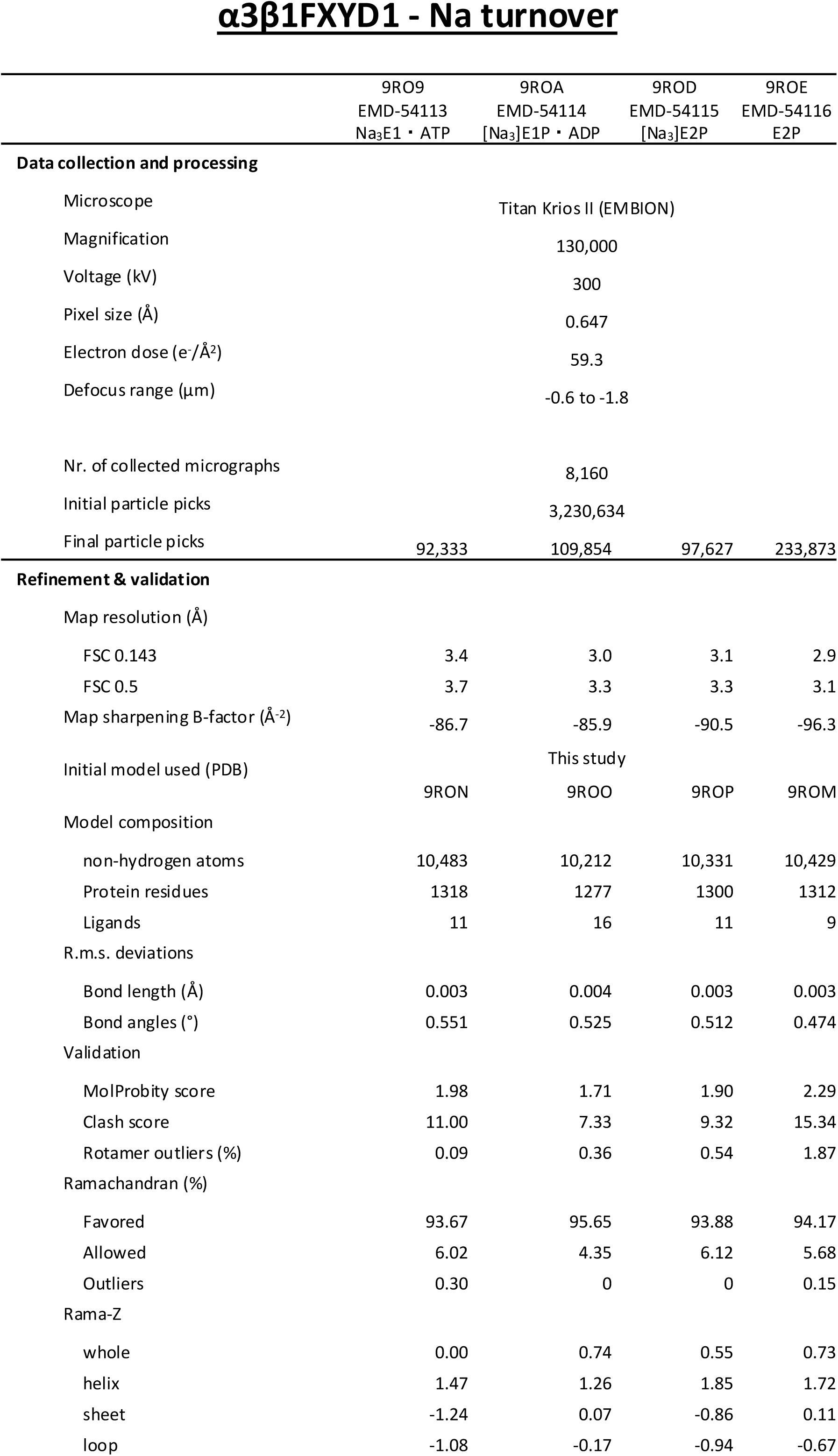
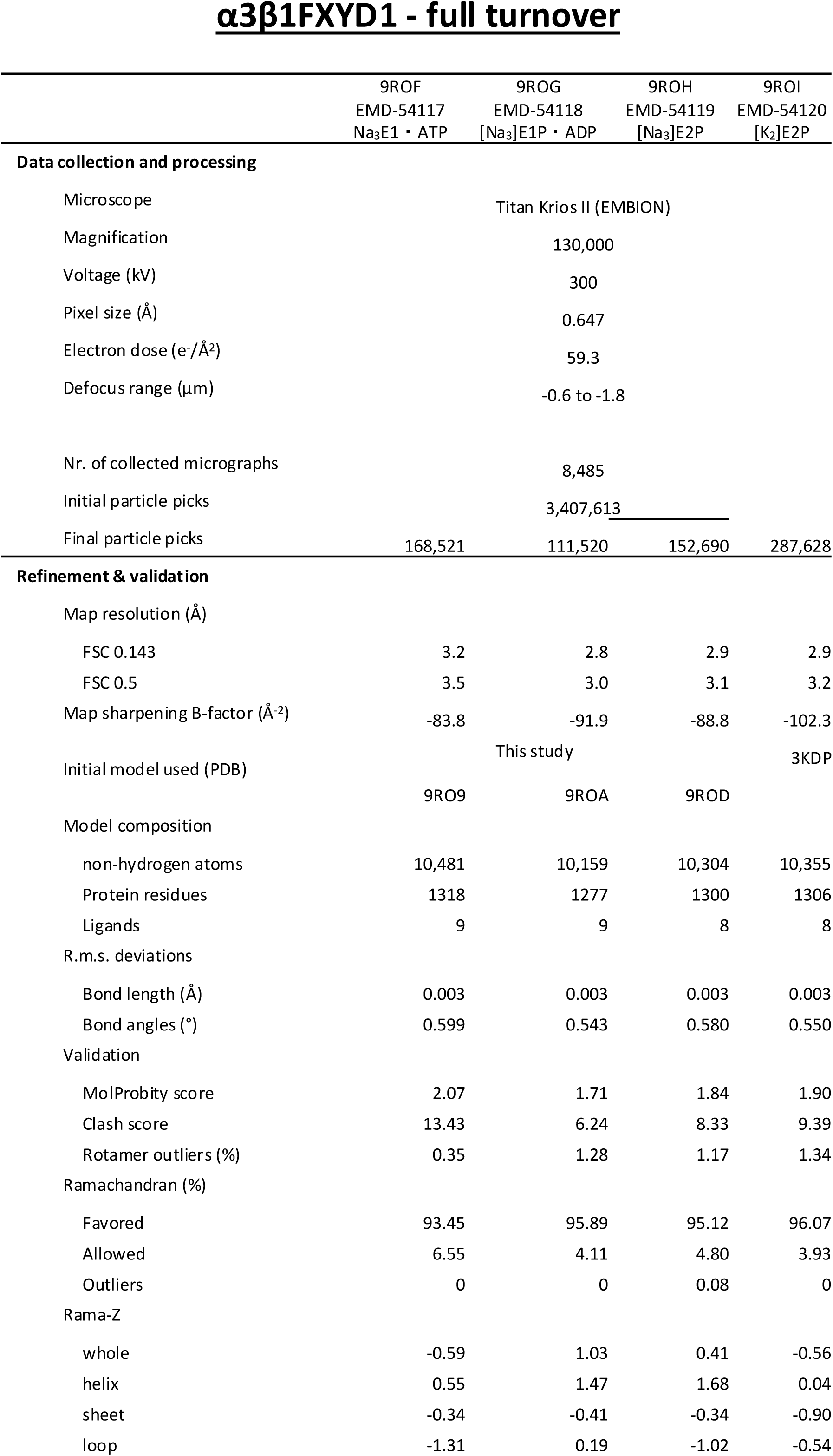
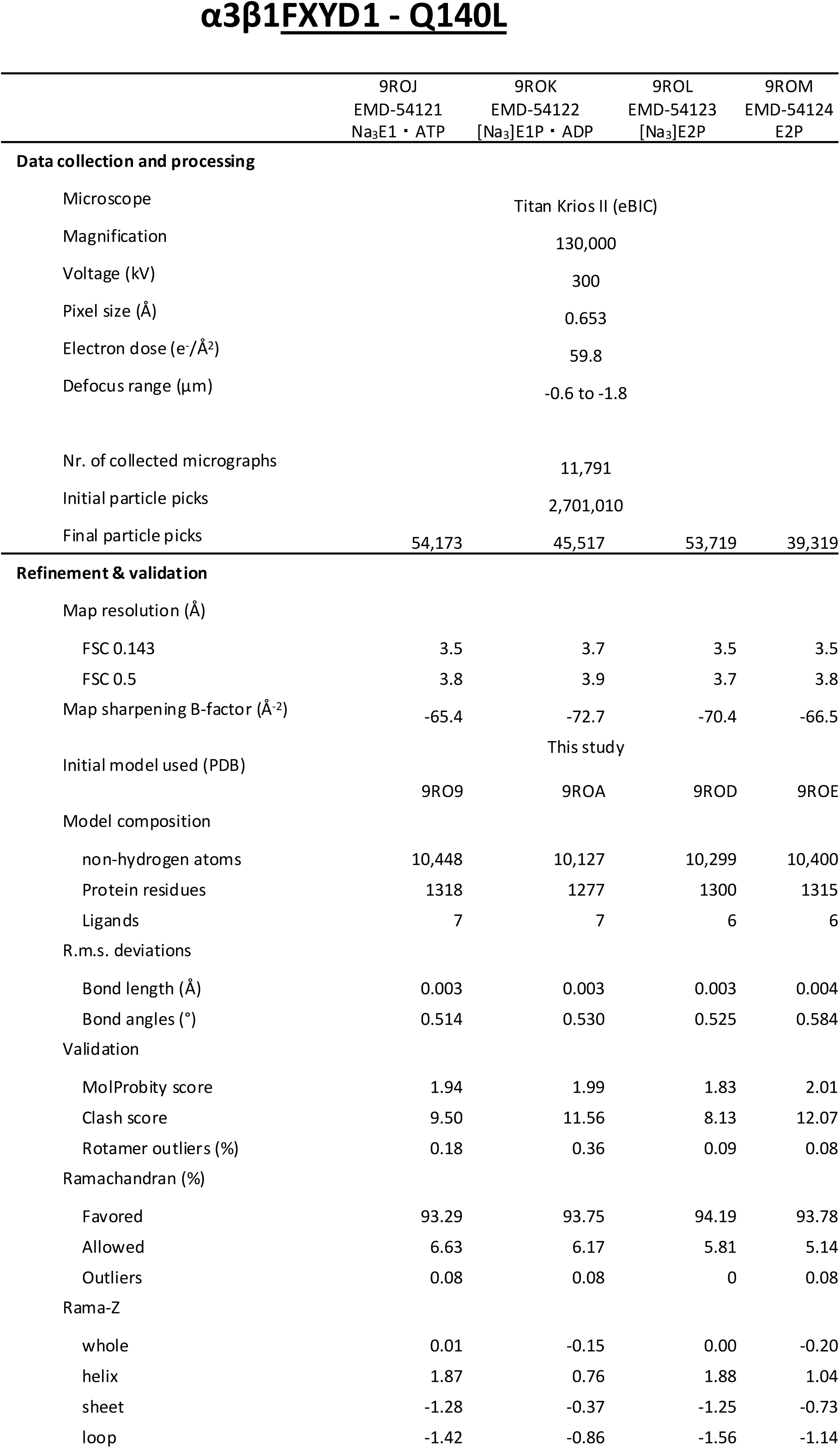
Cryo-EM data collection and processing, model building and refinement.

**Supplementary table 2.** Ouabain binding, Na,K-ATPase and turnover numbers determined in yeast membranes. Each value ±SEM represents the average of 3-5 determinations.

| Membranes | Ouabain binding (pmol/mg) | Na,K-ATPase activity (μmol/mg/min) | Turnover rate ($\text{min}^{-1}$) |
| --- | --- | --- | --- |
| $\alpha 1\beta 1$ (wt) | 15.5±1.4 | 0.13±0.006 | 8496±871 |
| $\alpha 1\text{Q150L}\beta 1$ | 10.2±0.85 | 0.019±0.004 | 1862±421 (22% of wt) |
| $\alpha 1\text{Q150H}\beta 1$ | 15.8±1.7 | 0.073±0.008 | 4620±709 (54% of wt) |
| $\alpha 3\beta 1$ (wt) | 8.0±0.5 | 0.044±0.0008 | 5500±358 |
| $\alpha 3\text{Q140L}\beta 1$ | 7.3±0.8 | 0.0038±0.001 | 520±148 (9.5% of wt) |

## Footnotes

1 An alternative indication for the relative poise of the E1/E2 equilibrium towards E2 in *α*3 comes from titrations of fluorescence quenching by Rb+ and reversal by Na+, which show a significantly higher apparent affinity of *α*3 for Rb^+^ compared to *α*1 (*α*3 K1/2 1.5±0.3mM versus *α*1 K1/2 4.3±0.44mM), coupled with a significantly lower apparent affinity for Na^+^ (*α*3 K1/2 26.7±1.2mM versus *α*1 K1/2 15.6 ±2.1 mM) (not shown).

